# Inhibiting mitochondrial translation overcomes multidrug resistance in MYC-driven neuroblastoma via OMA1-mediated integrated stress response

**DOI:** 10.1101/2023.02.24.529852

**Authors:** Karolina Borankova, Maria Krchniakova, Lionel YW Leck, Jakub Neradil, Adela Kubistova, Patric J Jansson, Michael D Hogarty, Jan Skoda

## Abstract

High-risk neuroblastoma remains a clinically challenging childhood tumor with a 5-year survival of only 50%. Tumors overexpressing N-MYC or c-MYC oncoproteins define a group of MYC-driven high-risk neuroblastoma with the most dismal outcomes, mainly caused by treatment failure due to the emergence and regrowth of multidrug-resistant cancer cells. Specific mitochondrial processes have been implicated in the maintenance of aggressive stem-like phenotypes in various cancers. We have recently identified a novel mitochondria-mediated mechanism of neuroblastoma multidrug resistance. However, the potential of pharmacological targeting of mitochondria to overcome therapy resistance and stemness in neuroblastoma remains unclear. Here, we show that c-MYC/N-MYC-driven multidrug-resistant neuroblastoma cells are highly vulnerable to cell death induced by the inhibition of mitochondrial translation. In contrast with normal fibroblasts, doxycycline (DOXY)-mediated inhibition of mitochondrial ribosomes efficiently impaired the survival of neuroblastoma cells regardless of their multidrug resistance and stem-like phenotypes. Mechanistically, inhibiting mitochondrial translation induced the mitochondrial stress-activated integrated stress response (ISR) via the OMA1-eIF2α axis, which preceded neuroblastoma cell death. Strikingly, several oncoproteins associated with poor neuroblastoma prognosis, including c-MYC and N-MYC, were markedly downregulated upon ISR activation. Comparing models of various neuroectodermal tumors and normal fibroblasts, we identified high levels of phosphorylated c-MYC and N-MYC (indicating their activity and rapid turnover) as a factor that predetermines susceptibility of neuroblastoma cells to DOXY-induced cell death. Neuroblastoma cells failed to develop significant DOXY resistance over a long-term repeated (pulsed) selection pressure, further demonstrating mitochondrial protein balance as a clinically relevant vulnerability of cancer cells that rely on high MYC activity. Together, our findings provide insight into mitochondrial retrograde regulatory networks in the context of MYC dependence and demonstrate the mitochondrial translation machinery as a promising therapeutic target in multidrug-resistant MYC-driven neuroblastoma.

## INTRODUCTION

Acquisition of aggressive dedifferentiated phenotype and therapy-induced multidrug resistance is the major cause of cancer therapy failure. Despite efforts, therapies that would overcome resistance mechanisms to kill all cancer cells including tumor-repopulating cancer stem-like cells remain elusive. Mitochondria have recently emerged as therapeutic targets in refractory cancers [1]. Besides serving as metabolic hubs, mitochondria integrate crucial roles in stemness maintenance [1], drug resistance [2], and cell death regulation [3]. Inhibiting mitochondrial processes has shown promising results in the most common tumor types, sensitizing resistant cancer cells to conventional chemotherapeutics [1]. However, our understanding of mitochondrial vulnerabilities in pediatric malignancies is limited.

Neuroblastoma is the most common extracranial childhood tumor. High-risk neuroblastomas are frequently driven by either N-MYC or c-MYC upregulation and have an extremely poor prognosis with 5-year overall survival of ~50% [4, 5]. Among other functions, MYC oncogenic transcription factors induce expression of genes involved in mitochondrial biogenesis [6–9] and mitochondria-dependent metabolism [10–12]. Importantly, we have recently demonstrated that mitochondria from therapy-resistant tumor cells often show attenuated apoptotic signaling, which largely contributes to neuroblastoma multidrug resistance [13]. Here, we therefore investigated potential mitochondrial dependencies, testing mitochondria as direct targets to overcome multidrug resistance in neuroblastoma. For this purpose, we took advantage of diverse mitochondrial inhibitors repurposed to target drug-resistant and/or stem-like cells in other cancers [14–18].

While mitochondria carry their own genome (mtDNA) and distinct transcription and translation machinery, their function heavily relies on nuclear-encoded mitochondrial proteins. Hence, mitochondrial perturbations must be efficiently relayed to cytosol and nucleus to orchestrate mitochondrial homeostasis, including proper stoichiometry of mitochondrial proteins [19]. Mitochondrial components of this retrograde signaling were only recently discovered, with inner mitochondrial membrane metalloprotease OMA1 identified as the most upstream regulator [20, 21]. We now provide insights into how this conserved mitochondrial stress-induced signaling might be exploited in neuroblastoma treatment. We demonstrate that disruption of mitochondrial proteostasis by inhibiting mitochondrial ribosomes activates OMA1-mediated integrated stress response (ISR), leading to c-MYC/N-MYC downregulation and cell death preferentially in neuroblastoma cells that rely on elevated MYC proteins, which offers a promising strategy to treat otherwise refractory MYC-driven tumors.

## RESULTS

### Mitochondria-targeting inhibitors overcome emergent multidrug resistance in post-therapy neuroblastoma cells

To establish a suitable model for our study, we first characterized a pair of *MFC*-amplified therapy-naïve CHLA-15 and drug-resistant CHLA-20 neuroblastoma cell lines derived from tumors of the same patient at diagnosis and at relapse after multimodal therapy, respectively [22]. Functionally, these cell lines differed in their stem cell-like characteristics, with CHLA-20 cells showing a markedly enhanced capacity to initiate tumors in NSG mice (**Fig. 1a)** and form neurospheres *in vitro* (**Fig. 1b**) while maintaining a similar growth rate (**Fig. 1c**). Pointing to a complex fine-tuning of stem-like traits [23, 24], both cell lines expressed similar levels of the c-MYC oncoprotein and other common stemness-associated markers, except for the upregulation of HIF-1α and OCT4 in CHLA-20 (**Fig. 1d, S1a**).

**Fig. 1:**
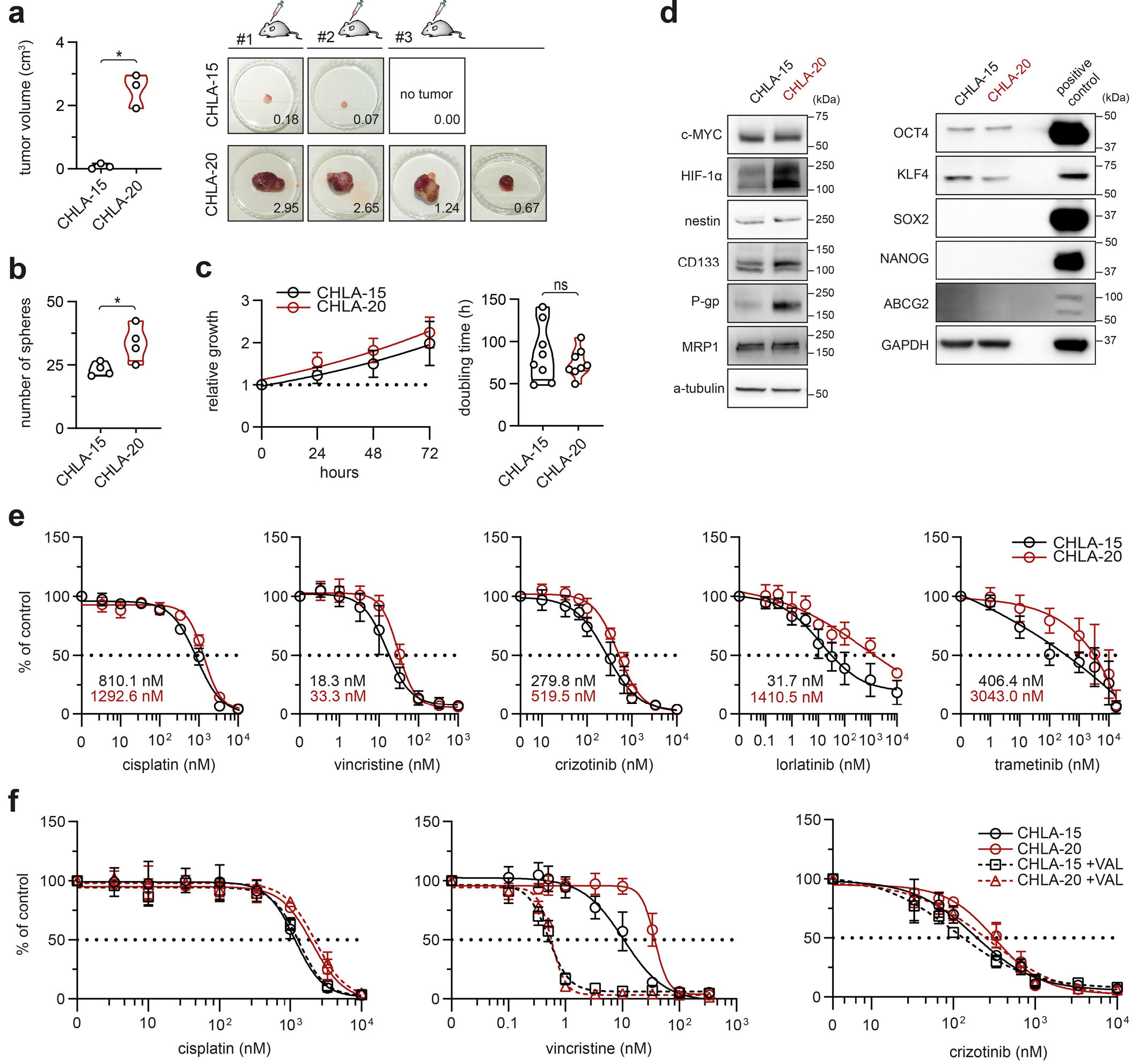
Therapy-naïve CHLA-15 and post-therapy CHLA-20 near-isogenic cell lines provide a useful model of therapy-induced drug resistance and cancer stemness in neuroblastoma. **a**, Mean volume of tumors per mice (left panel) and individual xenograft tumors (right panel) formed by CHLA-15 and CHLA-20 cells respectively after 29 days of injection in NSG mice. Right panel, numbers indicate the volume (cm^3^) of individual tumors. Also note the difference in the absolute tumor-forming efficiency (CHLA-15, 2/3 mice vs. CHLA-20, 3/3 mice). Left panel, data are presented as mean ± SD, biological n=3 mice per group. **b**, CHLA-20 are endowed with increased neurosphere formation capacity compared with CHLA-15, biological n=4, technical n=3. **c**, MTT cell viability assay analysis showed no significant difference in growth rate (left panel; data are mean ± SD) or calculated doubling times (right panel) of the cell line pair, biological n=8, technical n≥4. **d**, Western blot analysis of stemness transcription factors and CSC-related markers. Blots are representative of at least three experiments. Densitometric analysis is provided in Fig. S1. **e**, Sensitivity to diverse chemotherapeutics tested by MTT assay after 72 h of treatment (calculated IC_50_ values are indicated). Data are presented as mean ± SD, biological n=4, technical n=3. **f**, *In vitro* viability curves after 72 h exposure to drugs alone or with 0.5 μM of P-gp inhibitor valspodar (VAL). MTT data presented as mean ± SD, biological n=3, technical n=3. Statistical significance was determined by unpaired two-tailed Student’s t-test (**a-c)**, *p<0.05, ns – not significant.

Previously, CHLA-20 cells were reported more resistant to diverse chemotherapeutics than CHLA-15 [13, 22]. Validating this phenotype in our experimental settings, CHLA-20 cells exhibited broad resistance (1.6-fold to 44.5-fold difference vs. CHLA-15) to both DNA and non-DNA targeting drugs. The latter even included modern targeted agents never used to treat the donor patient’s tumor, i.e., crizotinib and lorlatinib, inhibiting ALK kinase (both CHLA-15 and CHLA-20 harbor the same *ALKR1275Q* activating mutation), or the MEK inhibitor trametinib (**Fig. 1e**). Of the common multidrug efflux pumps examined, only P-glycoprotein (P-gp) was upregulated in CHLA-20 compared with the therapy-naïve CHLA-15 (**Fig. 1d, S1b)**. However, P-gp upregulation was unlikely the mechanism underlying the broad resistance of CHLA-20 cells. First, it cannot explain resistance to cisplatin that is not a P-gp substrate. Second, when we inhibited P-gp by valspodar, CHLA-20 still retained enhanced resistance to drugs that are known P-gp substrates, such as crizotinib [25] (**Fig. 1f**). This is in line with our previous findings demonstrating that CHLA-20 has diminished sensitivity to apoptosis induction directly at the level of isolated mitochondria [13]. Together, the CHLA-15/CHLA-20 pair represents a useful model of emergent therapy resistance and cancer stemness in neuroblastoma.

To investigate potential mitochondrial vulnerabilities in this model, we treated both cell lines with several mitochondria-targeting drugs (**Fig. 2a,b**). Mitochondrial ATP production inhibitors, phenformin (inhibits complex I in the electron transport chain [1, 15]) and etomoxir (inhibits carnitine-palmitoyl transferase-1 [1, 14]), showed inefficient and reduced cell viability only at concentrations multiple times exceeding their selective or clinically tolerable profiles [26–29], with CHLA-20 still retaining a slightly increased resistance to these drugs (1.4-fold to 1.7-fold vs. CHLA-15; **Fig. 2b**). In contrast, inhibition of dynamin-related protein 1 (DRP1) by mdivi-1 and blocking mitochondrial protein synthesis by doxycycline (DOXY) substantially suppressed cell growth and induced apoptosis with a similar efficiency in both cell lines (**Fig. 2b-d, Supplementary Videos 1, 2**). Importantly, the effective DOXY concentrations (IC_50_ ~20 μM for 72 h) were within the range that shows a great clinical safety profile even during prolonged treatment [30, 31] and in young children [32, 33].

**Fig. 2:**
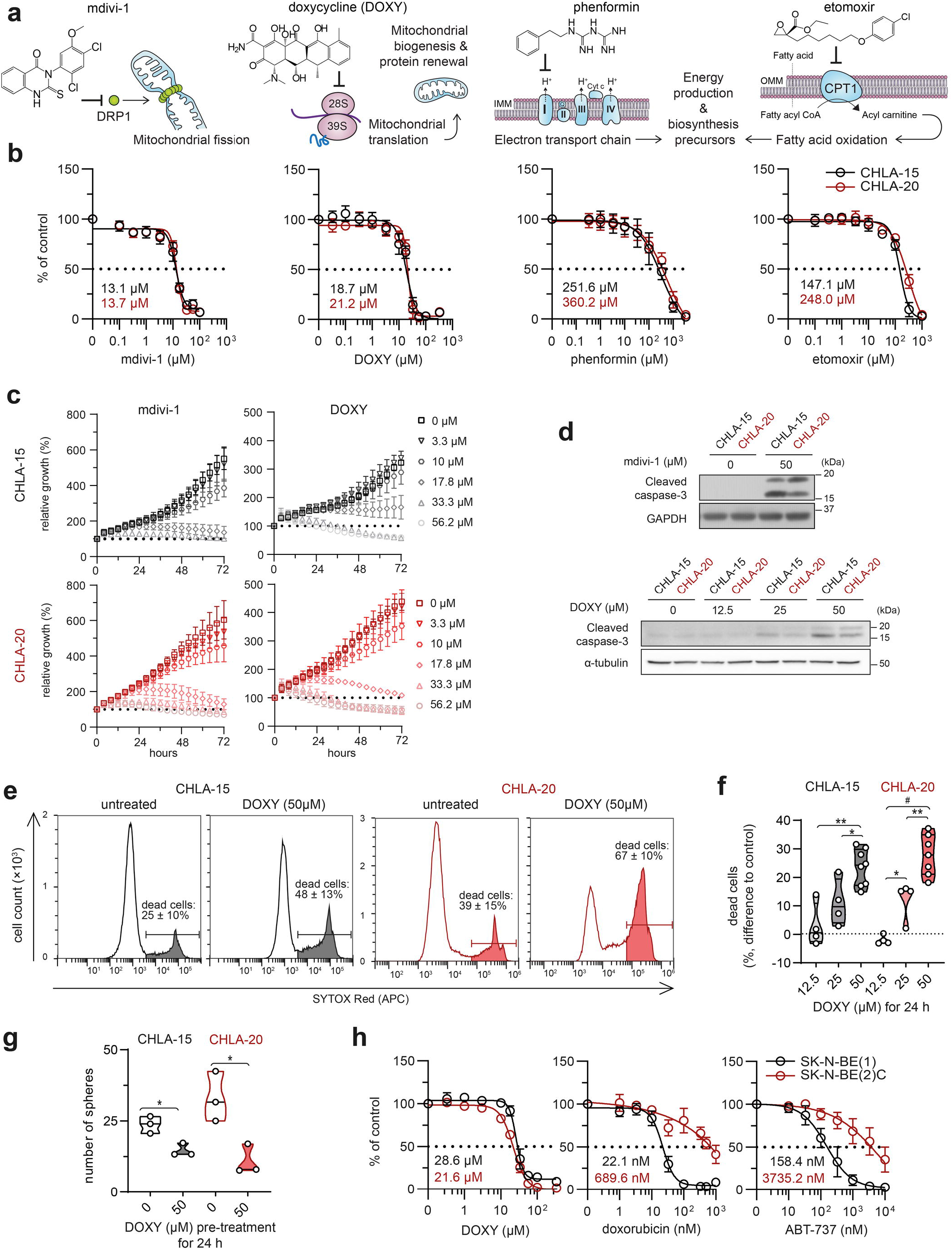
Therapy-naïve and drug-resistant neuroblastoma cells retain sensitivity to inhibitors of mitochondrial quality control. **a**, An overview of utilized mitochondrial inhibitors with indicated mechanism of action. **b**, MTT cell viability assay analysis after 72-h treatment showed no significant difference in sensitivity to mitochondrial inhibitors between therapy-naïve CHLA-15 and post-therapy CHLA-20 cell lines (calculated IC_50_ values are indicated). Data are presented as mean ± SD, biological n≥3, technical n=3. **c**, Live-cell imaging growth rate analysis of CHLA-15 and CHLA-20 treated with indicated concentrations of mdivi-1 and DOXY. Data are presented as mean ± SD, biological n≥3, technical n=3. Supporting Supplementary Videos 1,2 are provided. **d**, Western blotting of the cleaved caspase-3 showed that DOXY and mdivi-1 treatment for 24 h induced apoptosis in both CHLA15 and CHLA-20. Blots are representative of three experiments. **e,f**, Flow cytometry analysis of cell viability after DOXY treatment for 24 h. Representative histograms including percentages of SYTOX Red-positive dead cells as mean ± SD, biological n≥7 (**e**) and the difference in percentages of SYTOX Red-positive dead cells after indicated treatment vs. untreated controls (**f**). **g**, Pretreating neuroblastoma cells with 50 μM DOXY for 24 h reduced their neurosphere formation capacity. Notably, the inhibition of sphere-formation capacity was more pronounced in CHLA-20 cells which exhibit enhanced stem-like traits (3-fold reduction relative to untreated control) compared with CHLA-15 (1.6-fold reduction); biological n=3, technical n=3. **h**, CellTiter-Glo cell viability assay analysis of therapy-naïve SK-N-BE(1) and post-therapy SK-N-BE(2)C after 72-h treatment (calculated IC_50_ are indicated) showed their similar sensitivity to DOXY whereas post-therapy SK-N-BE(2)C were found resistant to conventional chemotherapy drugs (see also Fig. S2) or a BH3 mimetic, ABT-737, inhibiting multiple anti-apoptotic BCL-2 proteins. Data are presented as mean ± SD, biological n=3, technical n=3. Statistical significance was determined by one-way ANOVA followed by Tukey’s multiple comparisons test (**f**) and by unpaired two-tailed Student’s t-test (**g**), *p<0.05, **p<0.01, #p<0.0001.

Both mitochondrial translation and DRP1-mediated mitochondrial fission are crucial for mitochondrial renewal [34], suggesting that survival of neuroblastoma cells was dependent on efficient mitochondrial quality control irrespective of their drug-resistant state. We decided to explore this dependency by focusing on mitochondrial translation inhibition, using DOXY as one of the FDA-approved ribosome-targeting antibiotics that might be readily repurposed for potential anticancer therapies. Flow cytometric analysis confirmed the dose-dependent effects of DOXY leading to the induction of cell death in the CHLA-15/CHLA-20 pair (**Fig. 2e,f**). Viable DOXY-pretreated cells also formed significantly fewer spheres (**Fig. 2g**) which is a proxy for the capacity of DOXY to eliminate stem-like tumor-initiating neuroblastoma cells. We next validated the therapeutic potential of mitochondrial translation inhibition in another model pair of *MFCW*-amplified therapy-naïve SK-N-BE(1) and drug-resistant SK-N-BE(2)C neuroblastoma cells, the latter being even more sensitive to DOXY (**Fig. 2h, S2**). Collectively, these results revealed that targeting mitochondrial translation is efficient against bulk as well as drug-resistant/stem-like neuroblastoma cells.

### Inhibiting mitochondrial translation downregulates major oncoproteins and reveals a vulnerability shared across multiple neuroblastoma cell lines

Given the significant anticancer effects of DOXY, we further examined mechanisms underlying its activity. In line with previous studies [35–37], DOXY treatment induced an imbalance of mtDNA-encoded and nuclear-encoded mitochondrial proteins. While mtDNA-encoded cytochrome c oxidase I (MT-CO1) was downregulated in DOXY-treated cells, the levels of nuclear-encoded ATP synthase alpha-subunit 1 (ATP5A1) or mitochondrial import receptor subunit TOM20 homolog (TOMM20) were unaffected (**Fig. 3a,b**). Validating the general significance of our observations, we also detected similar growth inhibitory effects using other antibiotics that target bacterial, and thus mitochondrial ribosomes. Both tigecycline, a DOXY-related tetracycline derivate, and DOXY-unrelated antibiotics, linezolid and chloramphenicol, reduced cell viability in a dose-dependent manner (**Fig. 3c**). In contrast, ampicillin, a bacterial cell wall synthesis inhibitor not interfering with ribosome activity, did not affect the neuroblastoma cell growth (**Fig. 3d**). These results confirmed that the anti-neuroblastoma effects of DOXY were mediated by its specific binding to mitochondrial ribosomes leading to suppression of mitochondrial protein synthesis.

**Fig. 3:**
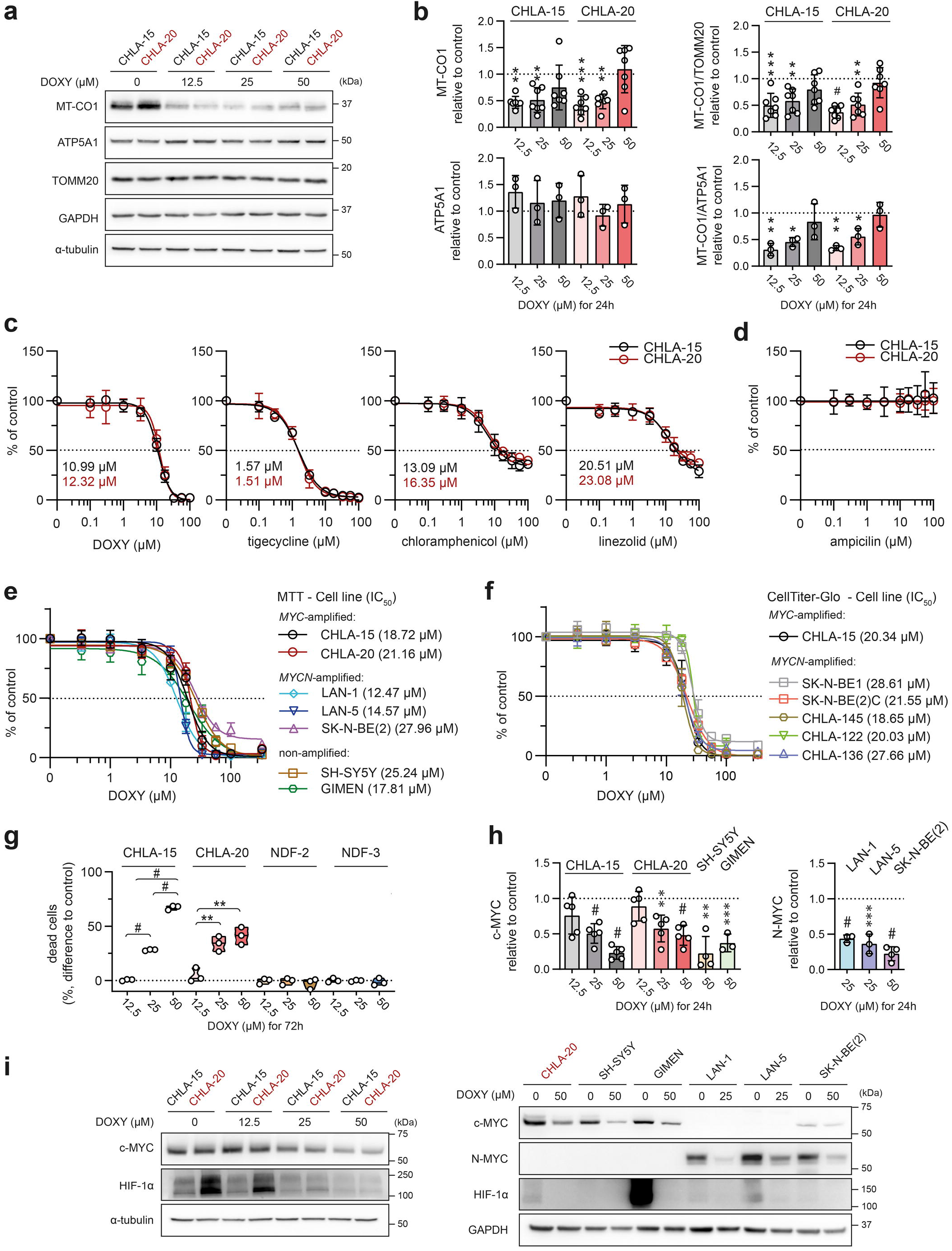
DOXY-mediated inhibition of mitochondrial protein synthesis impairs cell viability and reduces oncogenic transcription factors across a panel of neuroblastoma cells. **a,b**, Expression of mitochondrial proteins, mitochondrial-encoded MT-CO1 and nuclear-encoded proteins ATP5A1 and TOMM20, after 24-h DOXY treatment analyzed by western blotting (**a**) and subsequent densitometry (**b**). Normalized protein levels are plotted relative to untreated controls, mean ± SD. **c,d** MTT cell viability assay analysis of 6-day treatment with DOXY and other FDA-approved antibiotics targeting procaryotic ribosomes, tigecycline, chloramphenicol, and linezolid (**c**), and targeting bacterial cell wall synthesis, ampicillin **(d)**. Calculated IC_50_ are indicated. Data presented as mean ± SD, biological n=4, technical n=3. **(e)** MTT and **(f)** CellTiter-Glo cell viability assay analysis after 72-h treatment revealed all neuroblastoma cell lines to be highly sensitive to DOXY. Respective IC_50_ values are indicated in brackets. Data points are mean ± SD, biological n≥3, technical n=3. **g,** Cell death rate of neuroblastoma cells (CHLA-15 and CHLA-20) and neonatal dermal fibroblasts (NDF-2 and NDF-3) was analyzed after 72 h of DOXY treatment by flow cytometry using SYTOX Red staining. Data are presented as the difference in percentages of SYTOX Red-positive dead cells after indicated treatment vs. respective untreated controls, biological n=3. **h,i,** Densitometric analysis (**h**) of western blotting detection (**i**) revealed downregulation of c-MYC, N-MYC and HIF-1α across a panel of neuroblastoma cell lines treated with indicated concentrations of DOXY for 24 h. Normalized protein levels are plotted relative to untreated controls, mean ± SD. Densitometric analysis of HIF-1α is provided in Fig. S4. Statistical significance was determined by one-way ANOVA followed by Tukey’s multiple comparisons test (**b,g,h**), *p<0.05, **p<0.01, ***p<0.001, #p<0.0001.

We next utilized a panel of twelve neuroblastoma cell lines, including MYC- or MYCN-amplified and non-amplified clones, to test whether mitochondrial translation might be a common vulnerability in high-risk neuroblastoma. Using two different viability assays, we found all examined cell lines highly sensitive to DOXY (**Fig. 3e,f**). On the contrary, when tested in nonmalignant neonatal dermal fibroblasts NDF-2 and NDF-3, DOXY limited proliferation only at much higher concentrations and did not deteriorate cell viability (**Fig. 3g, S3, Supplementary Video 2)**, which suggests a potentially favorable therapeutic window of mitochondrial translation-targeted therapies in neuroblastoma.

Independent of genomic amplification, overexpression of N-MYC or c-MYC associates with the worst outcomes in neuroblastoma [38]. Yet, strategies targeting the MYC proteins for neuroblastoma treatment remain limited to preclinical studies [39]. Together with HIF-1α, another transcription factor associated with poor neuroblastoma prognosis, MYC proteins are known to affect mitochondrial biogenesis and metabolism [6–12]. Strikingly, DOXY-mediated inhibition of mitochondrial translation led to a dose-dependent downregulation of these transcription factors (**Fig. 3h,i, S4**), pointing to a novel therapeutically promising approach for inhibiting MYC proteins in multidrug-resistant high-risk neuroblastoma.

### DOXY disrupts mitochondrial morphology, suppresses mitochondrial fission machinery and primes mitochondria for apoptosis

Microscopically, most DOXY-treated neuroblastoma cells showed substantially impaired mitochondrial morphology and disrupted mitochondrial network (**Fig. 4a,b**). However, cleaved caspase-3 was detected only in a fraction of these cells, which suggests that the collapse of mitochondrial network was an early event after DOXY-induced inhibition of mitochondrial translation, priming mitochondria to apoptosis (**Fig. 4a**). As demonstrated by JC-1 probe, DOXY treatment disrupted mitochondrial membrane potential in a dose-dependent manner (**Fig. 4c**), which corresponded with the cleaved caspase-3 levels detected in cells treated with increasing concentrations of DOXY (**Fig. 2d**).

**Fig. 4:**
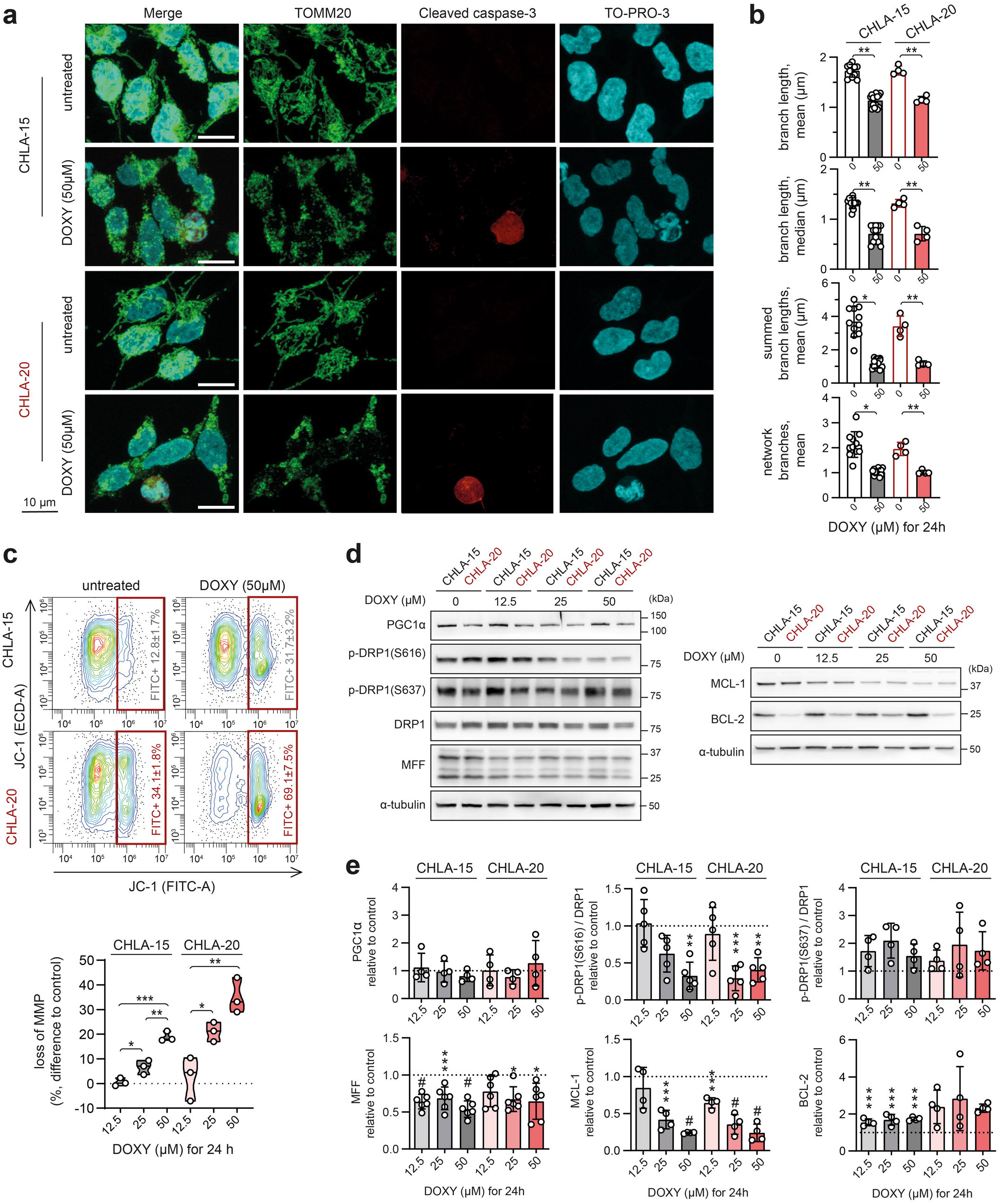
DOXY-mediated inhibition of mitochondrial translation disrupts mitochondrial morphology and fission machinery and primes mitochondria for apoptosis. **a,** Mitochondrial morphology–visualized by immunofluorescence staining of TOMM20 (green)–revealed signs of fragmentation and swelling in most cells after 24-h treatment with 50 μM DOXY. At this timepoint, apoptosis marked by cleaved caspase-3 (red) and fragmented nuclei (TO-PRO-3; cyan) was detected only in a subset of cells. Maximum intensity projections of confocal microscopy Z-stacks are shown. **b,** Image analysis by ImageJ plug-in tool MiNA - Mitochondrial Network Analysis confirmed disrupted mitochondrial morphology in cells treated with 50μM DOXY for 24 h. Data are presented as parameters determined for individual field of vision images, mean ± SD. **c,** Flow cytometry using JC-1 probe after 24-h DOXY treatment revealed dose-dependent loss of mitochondrial potential. Upper panel, representative contour plots; percentages of cells with depolarized mitochondria are presented as mean ± SD, biological n=3. Bottom panel, the differences in percentages of cells with depolarized mitochondria after indicated treatment vs. untreated cells. **d,e** Western blotting detection (**d**) and densitometric analysis (**e**) of proteins related to mitochondrial dynamics and BCL-2 anti-apoptotic proteins after 24-h DOXY treatment in indicated concentrations. Normalized protein levels are plotted relative to untreated controls, mean ± SD. Statistical significance was determined by unpaired two-tailed Student’s t-test (**b)** and by one-way ANOVA followed by Tukey’s multiple comparisons test (**e**), *p<0.05, **p<0.01, ***p<0.001, #p<0.0001.

The inhibition of mitochondrial translation did not affect the master regulator of mitochondrial biogenesis PGC1α (**Fig. 4d,e**). However, mitochondrial fission machinery, essential for the maintenance of mitochondrial health [40] and adjustment of mitochondrial functions [41], was suppressed after DOXY treatment. We found significant downregulation of fission-active DRP1 phosphorylated at Ser616, p-DRP1(S616), whereas its inactive form phosphorylated at Ser637, p-DRP1(S637), remained unaffected. Mitochondrial fission factor (MFF), a DRP1 adaptor protein, was also decreased upon DOXY treatment (**Fig. 4d,e**). At the level of anti-apoptotic BCL-2 family proteins, we identified a marked ~4-fold decrease of MCL-1 together with a ~2-fold BCL-2 upregulation (**Fig. 4d,e**). As CHLA-15 and CHLA-20 are both BCL-2-dependent [42, 43], these changes unlikely contributed to the DOXY-induced apoptosis.

### Mitochondrial stress-activated ISR is an early event during DOXY-mediated inhibition of mitochondrial translation

The identified mitochondrial protein imbalance, disrupted morphology and dynamics, and the loss of membrane potential collectively underpin a severe mitochondrial stress induced in DOXY-treated neuroblastoma cells. Recent studies showed that mitochondrial stress activates the OMA1-DELE1-HRI cascade, inducing the ISR by phosphorylation of its core mediator eIF2α [20, 21]. Due to the lack of reliable DELE1- or activated HRI kinase-specific antibodies, we assessed the activity of the key upstream and downstream regulators of this pathway. This analysis revealed consistent effects across a panel of neuroblastoma cells. Inhibiting mitochondrial translation activated mitochondrial stress sensor OMA1, marked by its autocatalytic depletion and cleavage of its substrate optic atrophy-1 (OPA1) [44], and led to the induction of ISR, marked by upregulation of phosphorylated eIF2α, p-eIF2α(S51), and the key ISR effector, CHOP [45] (**Fig. 5a,b, S5)**.

**Fig. 5:**
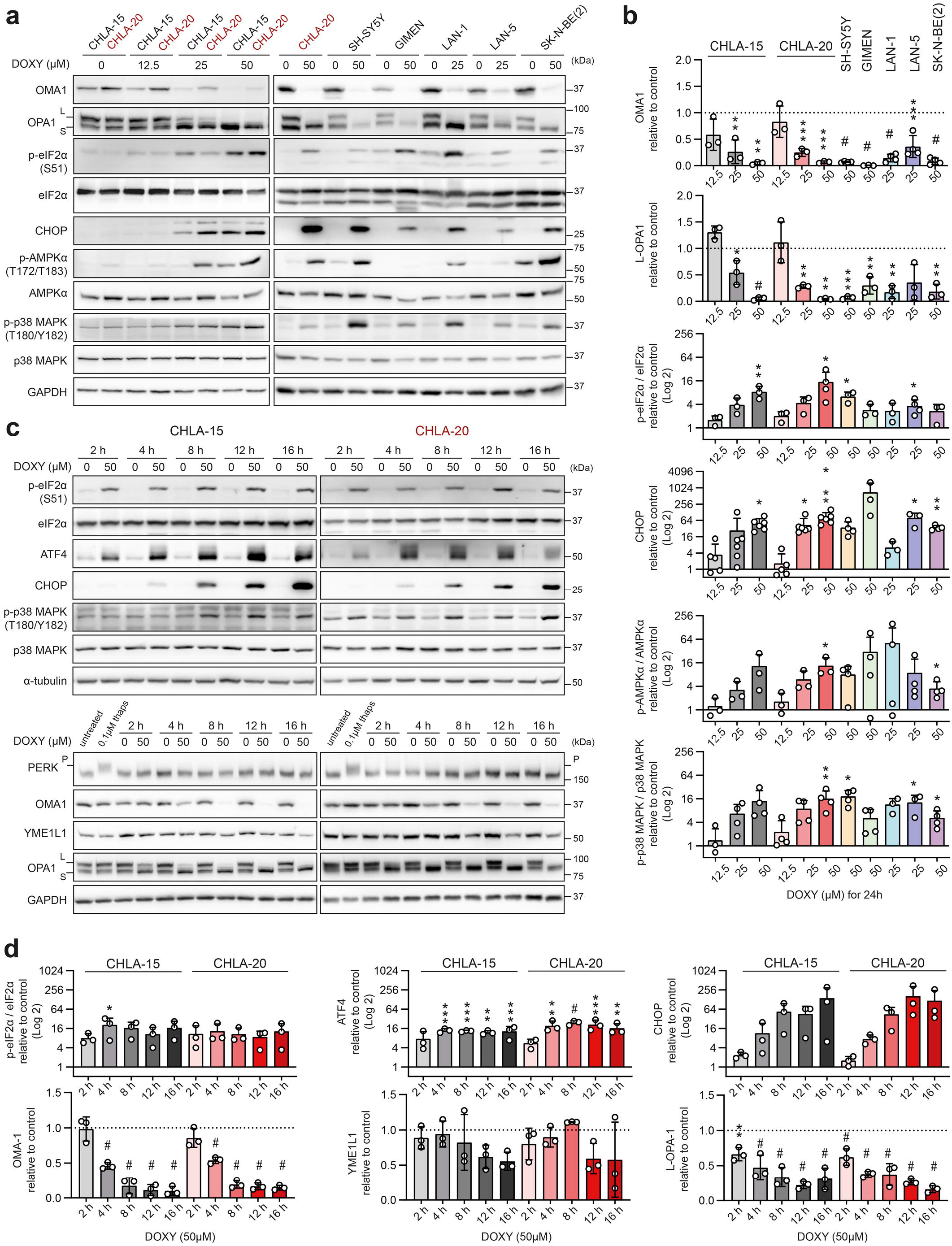
Inhibition of mitochondrial protein synthesis leads to early activation of OMA1-mediated ISR. **-a-d,** Western blotting detection and densitometric analysis of markers of mitochondrial stress, ISR and stress signaling kinases in a panel of neuroblastoma cells after DOXY treatment for 24 h **(a,b)** and over a 16-h time course with DOXY **(c,d)**. Normalized protein levels are plotted relative to untreated controls, mean ± SD. Treatment with 0.1 μM thapsigargin for 1 h served as a positive control of PERK phosphorylation, assessed by a reduced electrophoretic mobility in 6% polyacrylamide gel. Western blotting and densitometric analysis of YME1L1 after 24-h DOXY treatment and densitometric analysis of p-p38 MAPK (T180/Y182)/ p38 MAPK during 16-h DOXY treatment are provided in Fig. S7. Statistical significance was determined by one-way ANOVA followed by Tukey’s multiple comparisons test (**b,d**), *p<0.05, **p<0.01, ***p<0.001, #p<0.0001.

Long, intact L-OPA1 isoforms are crucial for the inner mitochondrial membrane fusion and their specific cleavage into short, fusion-inactive S-OPA1 forms by OMA1 induces the collapse of the mitochondrial network and promotes apoptosis [46, 47]. Consistently, the apparent degradation of L-OPA1 (**Fig. 5a-d**) likely explains the mitochondrial fragmentation observed in DOXY-treated cells, which counterintuitively showed impaired mitochondrial fission machinery (**Fig. 4**). Importantly, OMA1 activation (assessed by L-OPA1 processing) and increased levels of p-eIF2α(S51) and ATF4, a transcription factor promoting CHOP expression, were detected already after 2-h DOXY treatment (**Fig. 5c-d),** indicating the importance of mitochondrial ISR during the initial phase of mitochondrial translation inhibition.

PERK is a canonical kinase that phosphorylates eIF2α in response to endoplasmic reticulum (ER) stress from unfolded proteins. To evaluate whether ER stress was involved in the DOXY-induced ISR, we treated cells with ER stressor thapsigargin. However, PERK activation was detected only in thapsigargin-treated cells and, conversely, reciprocal degradation of mitochondrial stress-activated proteases OMA1 and YME1L1 and cleavage of L-OPA1 [44, 48] was found only in DOXY-treated samples (**Fig. 5c,d, S5, S6a-d**). These results demonstrate that DOXY-mediated inhibition of mitochondrial translation leads to mitochondrial stress that directly activates ISR independently of PERK and ER stress signaling.

Prolonged mitochondrial dysfunction is associated with ATP depletion, known to induce AMPK activity [49], and with overproduction of reactive oxygen species (ROS), activating redox-sensitive signaling cascades including p38 MAPK [50]. Indeed, we detected activation of these pathways, marked by increased levels of p-AMPKα(T172/T173) and p-p38 MAPK(T180/Y182), at later time points and DOXY concentrations that resulted in almost fully processed OPA1 and markedly degraded OMA1 (**Fig. 5a-d, S5**). Thus, the induction of AMPK and p38 MAPK signaling is a subsequent event following the mitochondrial stress-activated ISR in an OMA1-dependent fashion. However, we demonstrated that these late effects of DOXY-mediated inhibition of mitochondrial translation further contribute to neuroblastoma apoptosis, as inhibiting p38 MAPK by its specific inhibitor SB203580 partially prevented DOXY-induced activation of caspase-3 (**Fig. S6f,g**).

### Mitochondrial ISR links inhibited mitochondrial translation with degradation of short-lived oncoproteins

Phosphorylation of eIF2α attenuates cap-dependent translation in favor of ISR-specific mRNAs, allowing for efficient degradation of accumulated proteins and restoration of proteostasis [51]. We hypothesized that the ISR-mediated blockage of global protein synthesis might concurrently downregulate proteins with rapid turnover, as these are more readily targeted for degradation. Notably, c-MYC [52], HIF-1α [53] and MCL-1 [54], all significantly downregulated by DOXY treatment (**Fig. 3h,i**), are known to undergo rapid proteasome-dependent degradation with half-lives of <1 h. Indeed, the early mitochondrial stress-activated ISR was accompanied by the c-MYC, HIF-1α and MCL-1 downregulation that was detectable already upon 2-h treatment and progressed with time (**Fig. 6a,b**). In contrast, levels of BCL-2 that has a substantially longer half-life of ~20 h [55] were not significantly affected even after 16 h of DOXY treatment. Preferential downregulation of short-lived proteins upon the DOXY-induced ISR was further substantiated by the constant levels of the protein loading control GAPDH, which is a substrate for proteasome-independent chaperone-mediated autophagy with a half-life of ~40h [58] (**Fig. 6a,b**). In line with this reasoning, activation of the ISR by thapsigargin was also sufficient to downregulate c-MYC, HIF-1α and MCL-1 in neuroblastoma cells already within 1 h of treatment, whereas BCL-2 levels remained unchanged (**Fig. S6a,b**).

**Fig. 6:**
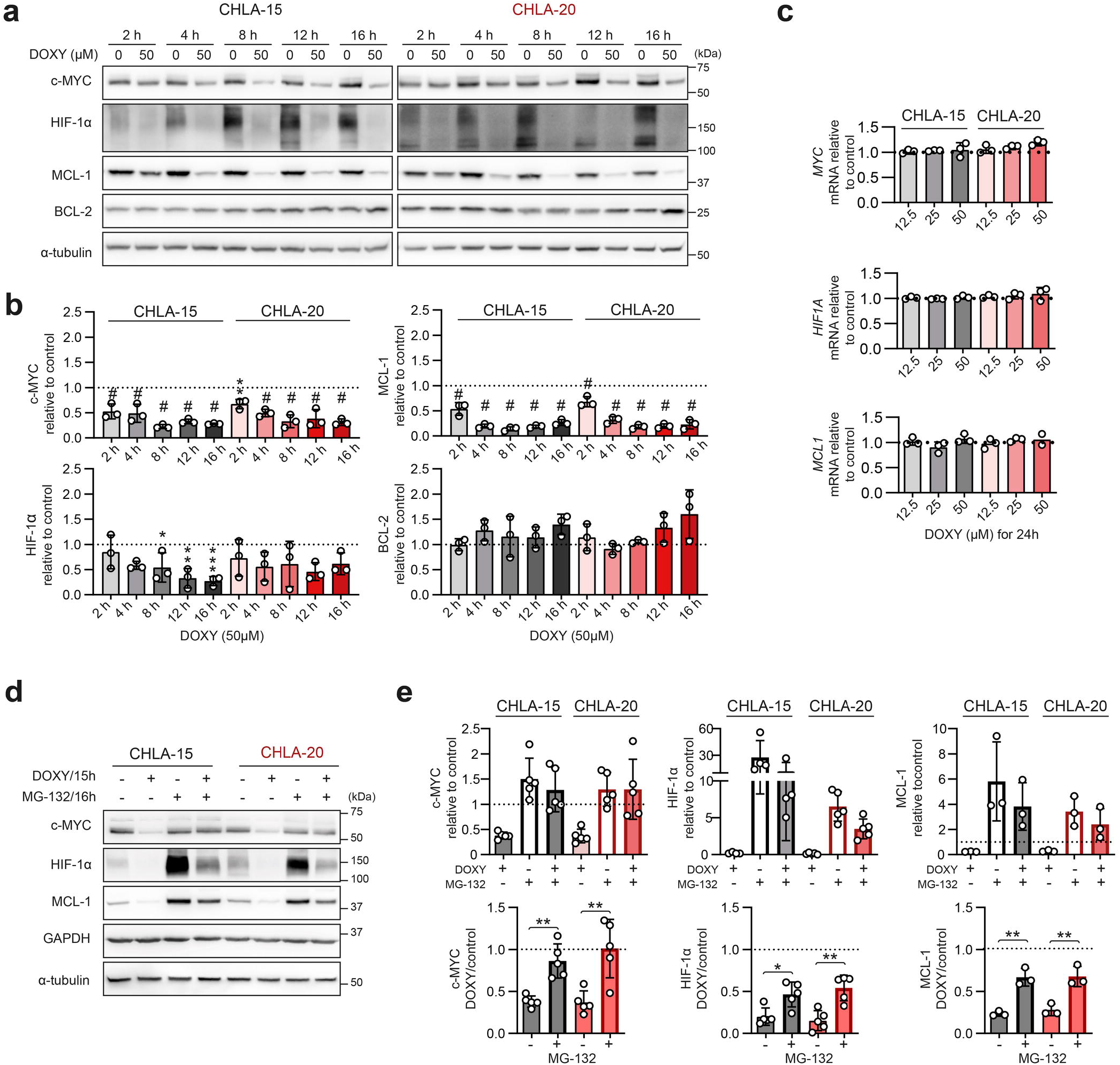
DOXY-induced downregulation of short-lived oncoproteins corresponds with early ISR activation and occurs exclusively at protein levels. **a,b,** Western blotting (**a**) and densitometric analysis (**b**) of c-MYC, HIF-1α, and anti-apoptotic BCL-2 family proteins in CHLA-15 and CHLA-20 treated with 50 μM DOXY for 2–16 h. Normalized protein levels are plotted relative to untreated controls, mean ± SD. **c,** Upon 24-h DOXY treatment analysis of mRNA levels of *MYC, HIF1A* and *MCL1* genes by RT-qPCR did not reveal any significant change on transcription level. Normalized mRNA levels plotted relative to untreated controls, mean ± SD. **d,e,** Western blotting detection (**d**) and densitometric analysis (**e**) of c-MYC, HIF-1α and MCL-1 after DOXY treatment of neuroblastoma cells with inhibited proteasome. To block proteasome activity, CHLA-15 and CHLA-20 were pre-treated with MG-132 (1 μM for MCL-1 analysis, 5 μM for c-MYC and HIF-1α) and after 1 h, 50 μM DOXY was added for additional 15 h. Upper panel, normalized protein levels are plotted relative to fully untreated controls (dashed line). Lower panel, ratios of normalized protein levels in DOXY-treated cells (w/o or with MG-132) and respective controls (w/o or with MG-132), mean ± SD. Statistical significance was determined by one-way ANOVA followed by Tukey’s multiple comparisons test (**b,c**) and unpaired two-tailed Student’s t-test (**e**), *p<0.05, **p<0.01, ***p<0.001, #p<0.0001.

We next confirmed that DOXY treatment did not affect expression of c-MYC, HIF-1α and MCL-1 at transcriptional levels (**Fig. 6c**). In contrast, proteasome inhibition by MG-132 almost completely reverted the DOXY-induced downregulation of c-MYC and significantly rescued expression of HIF-1α and MCL-1 (**Fig. 6d,e**). Thus, the attenuated protein synthesis and enhanced proteasomal degradation are the major mechanism causing the downregulation of short-lived oncoproteins upon mitochondrial stress-activated ISR in neuroblastoma cells.

### Rapid turnover of MYC proteins associates with sensitivity of MYC-driven neuroblastoma to DOXY-induced cell death

To explore if the effects of DOXY-mediated mitochondrial translation inhibition are neuroblastoma specific, we introduced seven cell lines derived from different nervous system tumors. Consistent with our results in neuroblastoma, inhibiting mitochondrial translation by DOXY limited cell proliferation across the tested cell lines, except for NSTS-5 schwannoma cells (**Fig. 7a**). Similarly, induction of mitochondrial stress, activation of ISR and decrease of HIF-1α were detected in DOXY-responsive glioblastoma, astrocytoma and medulloblastoma cells (**Fig. 7b-d, S7**). Intriguingly, markers of mitochondrial stress and ISR activation were also found in neonatal dermal fibroblasts (**Fig. S8)** that partially reduced proliferation but retained their viability in response to DOXY treatment (**Fig. 3g, S3)**. This may suggest that the ISR is a conserved retrograde pathway that relays DOXY-induced imbalance in mitochondrial proteins but leads to cell type-dependent outcomes.

**Fig. 7:**
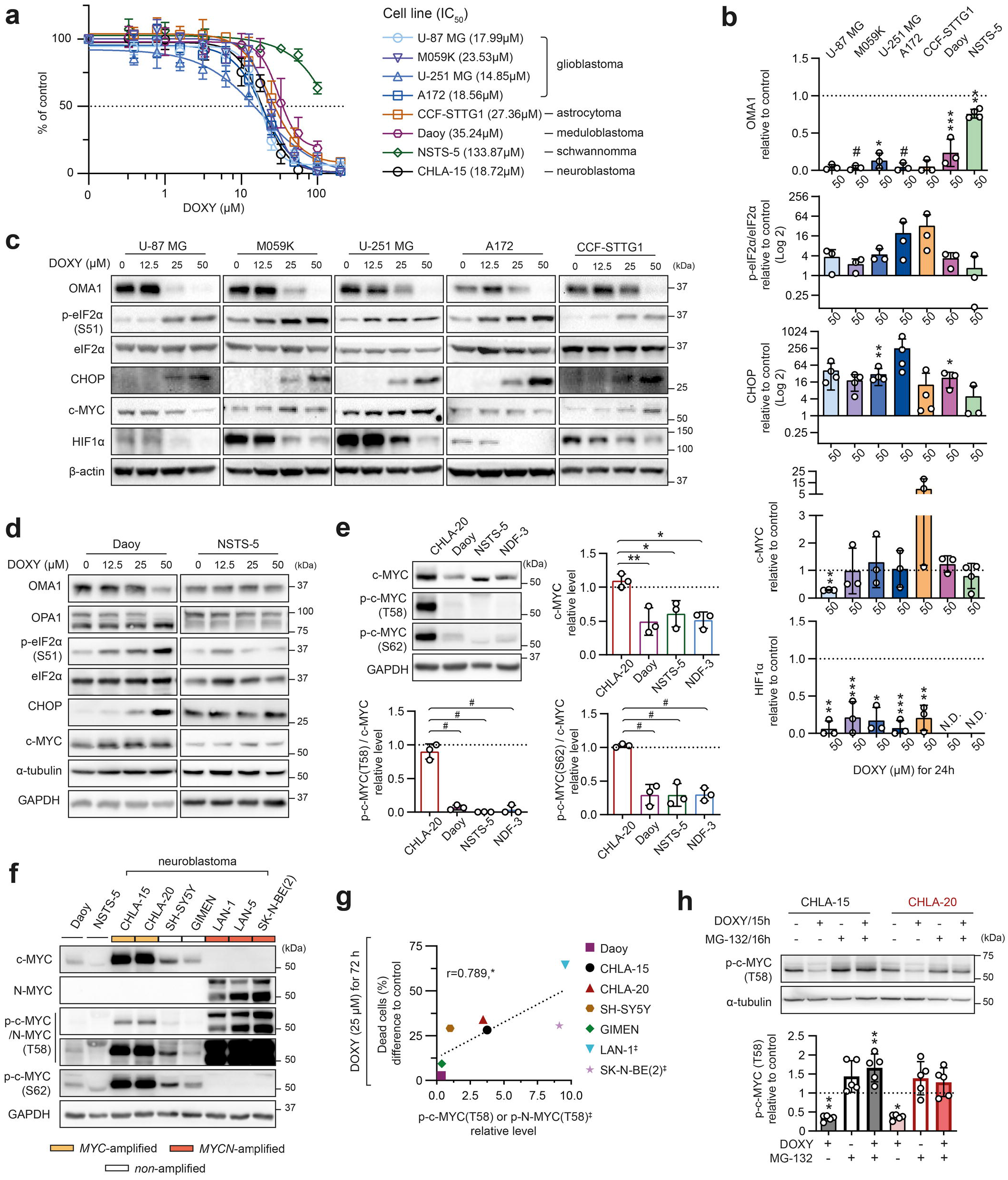
Overexpression and phosphorylation of MYC proteins correlate with sensitivity to cell death via mitochondrial ISR, conserved across various models of nervous system tumors. **a,** MTT cell viability assay analysis was used to determine sensitivity of cell lines derived from various nervous system tumors to DOXY. Respective IC_50_ values for 72-h treatment are indicated in brackets. Data presented as mean ± SD, biological n≥3, technical n=3. **b-d,** Densitometric analysis (**b**) of immunoblots (**c,d**) from lysates after 24-h DOXY treatment showed dose-dependent induction of mitochondrial stress and ISR in a panel of cell lines derived from different nervous system tumors. Contrary to neuroblastoma, this did not lead to consistent downregulation of c-MYC, although HIF-1α was significantly downregulated in nervous system tumors. Normalized protein levels are plotted relative to untreated controls, mean ± SD; N.D. – not detectable. Complete densitometric analysis is provided in Fig. S8. **e,** Western blotting detection and densitometric analysis of c-MYC, p-c-MYC(T58) and p-c-MYC(S62) revealed that CHLA-20 neuroblastoma cells have significantly higher c-MYC level and extensively enhanced phosphorylation of c-MYC compared with Daoy, NSTS-5 and NDF-3 cells that were less sensitive to DOXY. Normalized protein levels are plotted relative to untreated control, mean ± SD. **f,** Western blotting detection of c-MYC, p-CMYC/N-MYC(T58) and p-c-MYC(S62) in a panel of nervous system tumors. **g,** A significant positive correlation between the percentage of dead cells after 72-h treatment with 25 μM DOXY and the basal levels of p-CMYC/N-MYC(T58) in untreated neuroblastoma models and Daoy medulloblastoma cells, r = Pearson correlation coefficient; □, *MFCA*’-amplified. Related flow cytometry and protein densitometry data are provided in Fig. S10. **h**, Proteasome inhibition fully rescues DOXY-induced degradation of p-c-MYC(T58) in the CHLA-15/20 pair, as detected by western blotting (upper panel) and densitometric analysis (bottom panel). Cells were pre-treated with 5 μM MG-132 to block proteasome activity and after 1 h, 50 μM DOXY was added for additional 15 h. Normalized protein levels are plotted relative to fully untreated controls (dashed line). Statistical significance was determined by one-way ANOVA followed by Tukey’s multiple comparisons test (**b,e,h**) and by Pearson correlation (**g**), *p<0.05, **p<0.01, ***p<0.001, #p<0.0001.

In contrast to neuroblastoma, we did not observe consistent downregulation of c-MYC in cell lines derived from other nervous system tumors (**Fig. 7b-d, S7**) or neonatal dermal fibroblasts (**Fig. S8**). Compared with other tumor types, *MFC*-amplified CHLA-20 neuroblastoma cells express markedly higher levels of c-MYC together with its phosphorylated forms, p-c-MYC(T58) and p-c-MYC(S62) (**Fig. 7e**). Phosphorylation at threonine 58 (T58), subsequent to serine 62 (S62) phosphorylation, is known to promote proteasomal degradation and rapid turnover of both c-MYC [56] and N-MYC [57].

Strikingly, we found cell types with relatively low levels of c-MYC and its phosphorylated forms to be substantially resistant to DOXY-induced cell death, as demonstrated in Daoy (**Fig. S9**), NSTS-5 (**Fig. 7a**) and NDF-3 cells (**Fig. 3g, S3**). Immunoblots using the antibody recognizing both p-c-MYC(T58) and p-N-MYC(T58) showed that these phosphorylated forms are markedly upregulated in MYC-driven neuroblastoma and their levels correlated with sensitivity to DOXY-induced cell death in a panel of neuroblastoma and medulloblastoma models (**Fig. 7f,g, S10a,c,d**). Overall, our data indicate that MYC/MYCN status and the T58 phosphorylation, priming MYC proteins for rapid degradation upon the mitochondrial ISR (**Fig. 7h**), determined the propensity of MYC-driven neuroblastoma cells to cell death in response to inhibition of mitochondrial translation.

### Long-term repeated inhibition of mitochondrial translation does not induce highly resistant neuroblastoma phenotype

Target-specific resistance often develops between chemotherapy cycles. We therefore repeatedly exposed CHLA-15/CHLA-20 cells to DOXY to mimic multiple rounds of therapy and gradually select for the DOXY-resistant phenotype (DOXY-sel; **Fig. 8a**). However, even after 11 months and 30 cycles of selection, the cells remained highly sensitive to 2-fold of the initial IC_50_ of DOXY and the finally established DOXY-sel cell lines showed only ~1.5-fold resistance compared with parental cells **(Fig. 8b**). Both DOXY-sel cell lines exhibited downregulation of several proteins associated with poor neuroblastoma prognosis, including HIF-1α [58], P-gp, or MCL-1 (**Fig. S11a,b**). Given the marked P-gp downregulation in DOXY-sel cells, we next asked whether P-gp modulates sensitivity to DOXY. However, pharmacological inhibition of P-gp did not change the viability of parental cells treated with DOXY (**Fig. S11c**). Additional cell line-specific changes involved proteins targeted by DOXY treatment, including MT-CO1, eIF2α, or OMA1 (**Fig. S11a,b**). How these potentially compensatory events affect mitochondrial functions and the overall phenotype of DOXY-sel neuroblastoma cells will require further investigation.

**Fig. 8:**
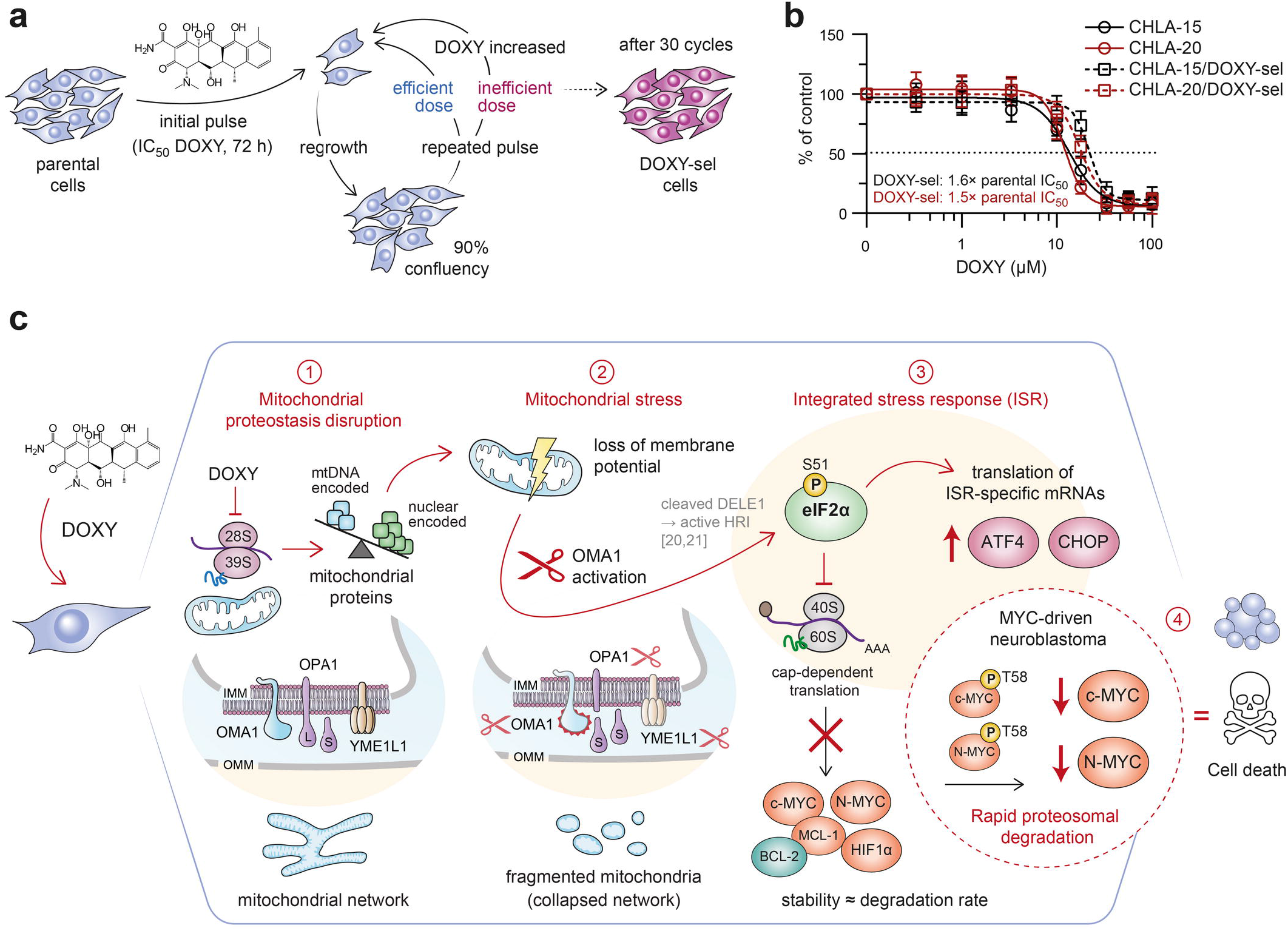
Inhibiting mitochondrial translation does not induce clinically relevant resistance and offers a promising therapeutic strategy for MYC-driven neuroblastoma. **a**, A schematic overview of the pulsed-selection strategy mimicking the chemotherapy cycles in the clinic. Parental cells CHLA-15 and CHLA-20 were repeatedly exposed to gradually increasing concentrations of DOXY, starting from IC_50_ (determined for each cell line by MTT assay upon 72-h treatment) as follows: 3 cycles of IC_50_, followed by 3 cycles of 1.5-fold IC_50_ and finally 2-fold IC_50_ that remained highly efficient for additional 24 cycles. In each cycle, cells were treated for 72 h and let to regrow to approx. 90% confluency in fresh drug-free culture media before another round of DOXY treatment. **b**, The sensitivity of parental cells and established DOXY-selected cells (CHLA-15/DOXY-sel and CHLA-20/DOXY-sel) to DOXY was compared by MTT cell viability assay after 72 h of treatment. Data are presented as mean ± SD, biological n=3, technical n=3. **c**, A model summarizing synthetic lethal effects induced by inhibition of mitochondrial translation in MYC-driven neuroblastoma. Blocking mitochondrial ribosomes by DOXY disrupts mitochondrial proteostasis (1) which impairs mitochondrial function and leads to the mitochondrial stress-induced activation of OMA1 (2). Activated OMA1 mediates OPA1 cleavage, eventually resulting in collapsed mitochondrial network, and relays mitochondrial stress directly via the mitochondrial ISR without involving ER stress signaling (3). Besides preferential translation of ISR-specific mRNAs, attenuated cap-dependent translation leads to downregulation of short-lived proteins primed for rapid degradation, including p-c-MYC(T58) and p-N-MYC(T58), which sensitizes MYC-driven neuroblastoma to cell death induced by the inhibition of mitochondrial translation (4).

Collectively, our data establish that perturbing mitochondrial function by targeting mitochondrial protein synthesis triggers the mitochondrial ISR which in the context of MYC-driven neuroblastoma efficiently overcomes existing multidrug resistance mechanisms and eradicates aggressive tumor cells without inducing clinically relevant resistance (**Fig. 8c**).

## DISCUSSION

Targeting mitochondria is an emerging strategy to overcome cancer drug resistance. Yet, molecular determinants that would guide efficient mitochondrial therapies are poorly understood, particularly in pediatric tumors. Here, we identified mitochondrial translation as a promising MYC-dependent target in multidrug-resistant neuroblastoma. Our results also implicate that mitochondrial quality mechanisms, including mitochondrial dynamics and translation control, are essential for neuroblastoma cell survival irrespective of the drug resistance status, as demonstrated by inhibition of mitochondrial fission by mdivi-1 and disruption of mitochondrial ribosome processivity by DOXY (**Fig. 2c-h, Supplementary Videos 1,2**).

Mechanistically, time course experiments showed that inhibition of mitochondrial ribosomes by DOXY induces the activity of mitochondrial metalloprotease OMA1 and initiates mitochondrial retrograde signaling via the central ISR mediator eIF2α. Short-running ISR normally orchestrates gene expression and protein translation to restore cellular homeostasis and maintain cell survival, in line with its reported pro-tumorigenic effects [59]. However, excessive or prolonged ISR is known to induce cell death in various cell types [51]. Interestingly, the ISR effector ATF4 was already shown to mediate cell death upon glutamine deprivation in *MFCV*-amplified neuroblastoma [60]. Consistently, we provide evidence that MYC-driven neuroblastoma cells are particularly vulnerable to the ISR-mediated cell death, while p38 MAPK appears to further favor apoptosis induced by the mitochondrial ISR. p38 MAPK is known to promote apoptosis [61–63] and likely synergizes with the ISR by directly phosphorylating CHOP [64], enhancing its pro-apoptotic activity [65].

Previously, DOXY-mediated inhibition of mitochondrial translation was shown to enhance ER-mitochondria connectivity [66] and was suggested to activate ISR via inducing ER stress in carcinomas [66, 67] and melanoma [18, 68]. On the contrary, we demonstrate that ER stress is not involved in DOXY-induced ISR in neuroblastoma, as eIF2α phosphorylation was not mediated by its upstream ER stress-activated kinase PERK. Our data are consistent with recent studies demonstrating mitochondrial stress sensor OMA1 as the major inducer of ISR via the OMA1-DELE1-HRI pathway [20, 21]. Similarly, we demonstrate that DOXY-induced mitochondrial protein imbalance disrupts mitochondrial morphology and membrane potential, inducing the ISR along the OMA1 activity detected already in early phases of the treatment. These results strongly support the OMA1-dependent activation of ISR upon blocking mitochondrial ribosomes in neuroblastoma, but they do not exclude additional cell context-dependent molecular regulators. Previously, knocking down components of the OMA1-DELE1-HRI pathway showed insufficient to completely block the DOXY-mediated ISR in embryonal kidney HEK293T cells [20]. However, in contrast to our models (**Fig. 5, 7d, S5a,b**), DOXY-treated HEK293T cells did not show any signs of L-OPA1 cleavage [20], suggesting cell type-dependent activation of OMA1 in response to the inhibition of mitochondrial translation.

A chronic ISR has been recently suggested to predict the sensitivity of melanoma cells to tigecycline-mediated inhibition of mitochondrial ribosomes [18]. In contrast, we did not observe any association between DOXY-induced cell death and the extent of ISR activation prior or after treatment. In fact, we found p-eIF2α(S51) and one of the major ISR effectors, CHOP, to be consistently upregulated by DOXY in nearly all tested cell lines (neuroblastoma: 11/11; other nerve tissue tumors: 6/7; normal fibroblasts: 1/1) regardless their vulnerability to DOXY-induced cell death. High c-MYC levels have been associated with the susceptibility of hematological malignancies to inhibition of mitochondrial protein synthesis [37, 69]. In this study, we found that the DOXY-induced mitochondrial ISR leads to degradation of short-lived MYC proteins and that the upregulation of their T58-phosphorylated forms, tagging MYC proteins for rapid proteasome degradation [56, 57, 70, 71], likely sensitizes MYC-driven neuroblastoma to inhibition of mitochondrial ribosomes.

Based on these findings, we propose mitochondrial translation as a novel synthetic lethal target that might be exploited to overcome multidrug resistance in MYC-driven neuroblastoma (**Fig. 8c**). We anticipate that further dissection of OMA1-regulated pathways might enable identification of additional clinically relevant targets for the treatment of MYC-addicted tumors.

## METHODS

### Cell culture and treatment

The following cell lines were used in the study: (i) neuroblastoma cell lines CHLA-15, CHLA-20, CHLA-122, CHLA-136, CHLA-145, SK-N-BE(1), SK-N-BE(2)C (obtained from the COG/ALSF Childhood Cancer Repository; www.cccells.org), GIMEN, LAN-1, LAN-5 (a kind gift of Prof. Lumír Krejčí), SH-SY5Y, and SK-N-BE(2) (purchased from ECACC); (ii) glioblastoma cell lines U-87 MG, M059K, U-251 MG, A-172 and astrocytoma cell line CCF-STTG1 (all purchased from ATCC); (iii) medulloblastoma cell line Daoy (purchased from ECACC); (iv) schwannoma cell line NSTS-5 (in-house derived from tumor tissue with written informed consent under IGA MZCR NR/9125-4 project approved by the Research Ethics Committee of the School of Medicine, Masaryk University, Brno, Czech Republic – approval no. 23/2005); (v) human neonatal dermal fibroblast cell lines NDF-2 and NDF-3 (#CC-2509; Lonza Bioscience, Durham, NC, USA; a kind gift of Dr. Tomáš Bárta). In addition, human pluripotent embryonal carcinoma cell line NTERA-2 (clone D1) purchased from ECACC (#01071221) served as a positive control shown in Fig. 1d. All cell lines were authenticated by STR profiling (Generi Biotech, Hradec Králové, Czech Republic; Westmead Institute of Medical Research, Westmead, NSW, Australia; Promega Geneprint 10, Madison, Wisconsin, USA) and routinely tested for mycoplasma by PCR [72].

Cell lines were cultured in a humidified atmosphere of 5% CO_2_ at 37°C in media with supplements as detailed in Supplementary Table 1,2. Drug treatments were always performed the day after cell seeding. A detailed overview of the drugs used is provided in Supplementary Table 3.

### Cell viability assays

For 24–72-h and 6-day experiments, cells were seeded into 96-well plates in a density of 5000 and 2500 cells/well, respectively. After the incubation time, cell viability was measured. Performing MTT assay, thiazolyl blue tetrazolium bromide (#M2128, Sigma-Aldrich, St. Louis, MO, USA) was added to each well to reach the final concentration of 0.455 mg/ml. After 3-h incubation under standard conditions, the medium was replaced with 200 μL of DMSO to solubilize the formazan crystals. The absorbance of each well was determined by Sunrise Absorbance Reader (Tecan, Männedorf, Switzerland). For CellTiter-Glo® assay (#G7571, Promega, Madison, WI, USA), cells were seeded in Corning® 96-well Flat Clear Bottom White Polystyrene TC-treated luminescent microplates (#3610, Corning, Corning, NY, USA) and manufacturer instructions were followed. Luminescence was measured by Synergy™ 2 microplate reader (BioTek Instruments, Winooski, VT, US).

Absolute half maximal inhibitory concentration (IC_50_) of drugs and inhibitors was determined from non-linear regression of datasets of MTT assay and CellTiter-Glo® assay with individual tested concentrations normalized to untreated control cells. Non-linear regression with variable slope was calculated using GraphPad Prism 8.0.2. software (GraphPad Software, San Diego, CA, US). Determined parameters were then used to calculate absolute IC_50_ values according to the following formula: relative IC_50_*(((50-top)/(bottom-50))^(−1/hill slope)).

### Growth analysis

5000 cells/well were seeded into 96-well plates and treated the day after seeding with respective drugs. Subsequently, cell confluency was determined every 4 hours using live cell imaging system Incucyte® S3 (Sartorius, Göttingen, Germany) and plotted relative to initial values.

### Sphere formation assay

Cells were harvested and dissociated into single-cell suspension using Accutase (#LM-T1735, Biosera) and seeded into ultra-low attachment 6-well plates (#CLS3471-24EA, Corning) at a density of 1000 cells/well in a defined serum-free medium: DMEM/F12 based (as detailed in Supplementary Table 1) w/o fetal bovine serum, supplemented with 1× B27 w/o vitamin A (#12587, Gibco), 10 ng/ml EGF (#E9644, Sigma-Aldrich), and 20 ng/ml FGF2 (#SRP4037, Sigma-Aldrich). Cells were replenished with 200 μl of the freshly prepared medium three times a week. The number of spheres (diameter ≥ 50 μm) was manually counted after 21 days of incubation using PROView software analysis of images taken under IM-3 light microscope equipped with C-B5 digital camera (all Optika, Ponteranica, Italy).

### Tumorigenicity assay *in vivo*

Cells were harvested, enzymatically dissociated and a single cell suspension of 1□×□10^6^ cells in 100 μl of pure DMEM/F/12 medium (#LM-D-1224, Biosera) was injected subcutaneously into the right flank of nine-week-old female NSG (NOD/ShiLtSz-*scid/I12rγ*^null^) mice. All animal experiments were conducted in accordance with a study (MSMT-4408/2016-6) approved by the Institutional Animal Care and Use Committee of Masaryk University and registered by the Ministry of Education, Youth and Sports of the Czech Republic as required by national legislation. After 29 days, the mice were sacrificed and surgically examined. The xenograft tumors were excised and photographed, and the final tumor volume was determined using the following formula: tumor volume (mm^3^) = length (mm)□×□width (mm)□×□width (mm)□×□1/2.

### Western blotting

Whole-cell extracts were collected using RIPA lysis buffer (2 mM EDTA, 1% IGEPAL® CA-630, 0.1% SDS, 8.7 mg/ml sodium chloride, 5 mg/ml sodium deoxycholate, 50 mM Tris-HCl) supplemented with cOmplete™ Mini Protease Inhibitor Cocktail (#11836170001, Roche, Basel, Switzerland) and PhosSTOP (#4906837001, Roche). 20 μg of total proteins were resolved on 10% polyacrylamide gels (except for 6% gels used for PERK detection) and blotted onto PVDF membranes (#1620177, Bio-Rad Laboratories, Hercules, CA, USA). The membranes were blocked with 5% not-fat dry milk or bovine serum albumin (#A7906, Sigma-Aldrich) in Tris-buffered saline with 0.05% Tween-20 (#93773, Sigma-Aldrich) for at least 1 h and incubated with primary antibodies overnight on rocking platform at 4°C. The incubation with secondary HRP-linked antibody was conducted at RT for at least 1 h. The list of antibodies used, including dilutions and respective blocking agents, is provided in Supplementary Table 4. Chemiluminescent detection was performed following a 5-min incubation with ECL™ Prime Western Blotting Detection Reagent (#RPN2236, Cytiva, Marlborough, MA, USA) using either Azure C600 imaging system (Azure Biosystems, Dublin, CA, USA) or light sensitive films (#CP-BU NEW 100 NIF, Agfa, Mortsel, Belgium).

Densitometric analysis of western blotting images was done using gel analysis tool in ImageJ (Fiji) software (NIH, Bethesda, MD, USA), version 2.1.0/1.53c. The signal of a protein of interest was normalized to that of a loading control, α-tubulin, β-actin, or GAPDH, detected on the same gel. For all relevant figures, original uncropped blots including all replicates are provided as supplementary Original Data file.

### Immunostaining

Cells were seeded on coverslips coated with Matrigel (#734-1440, Corning). After the incubation period, the coverslips were rinsed by PBS and cells were fixed by 3% paraformaldehyde (#158127, Sigma-Aldrich). The cells were then permeabilized using 0.2% Triton X-100 (#04807423, MP Biomedicals, Irvine, CA, USA) for 1 min and blocking was performed by 3% bovine serum albumin (#A7906, Sigma-Aldrich) for 10 min at RT. The primary and secondary antibodies (Supplementary Table 4) were diluted in blocking solution. The incubation with antibodies lasted for at least 60 minutes. Nuclei were stained by TO-PRO3 (#T3605, Invitrogen, Carlsbad, CA, USA). The cells were mounted by ProLong™ Diamond Antifade (#P36961, Invitrogen) and imaged using Leica SP8 confocal microscope (Leica, Wetzlar, Germany). Z-stacked images were captured and processed as maximum intensity projections using software LAS X (Leica, 3.4.218368).

### Mitochondrial morphology analysis

Mitochondrial morphology was determined from maximum intensity projection of z-stack confocal images of TOMM20 fluorescence channel using ImageJ (Fiji) plug-in tool MiNA (Mitochondrial Network Analysis) [73]. The pre-processing parameters were applied as follows: 3rd-order median filter with radius 1; 1st-order unsharp mask with radius 2 and mask weight 0.9; contrast enhancement using 2nd-order CLAHE with block size 127 pixels, histogram bins 256 and maximum slope 3.

### Flow cytometry

For all experiments, cells were harvested using Accutase (#LM-T1735, Biosera) and diluted in PBS with 3% fetal bovine serum (#FB-1101, Biosera) and 2mM EDTA (#ED2SS, Sigma-Aldrich) and immediately processed for measurements using CytoFLEX S flow cytometer (Beckman Coulter, Brea, CA, USA). Cell viability was measured following 15-minute incubation with 5 nM SYTOX™ Red Dead Cell Stain (#S34859, Invitrogen) on ice as per manufacturer’s instructions.

Mitochondrial membrane potential was assessed by 10 μg/ml JC-1 Probe (#65-0851-38, Invitrogen) after 10-min incubation at 37°C. Fluorescence of JC-1 monomers and aggregates was excited by 488 nm laser and detected using FITC (525/40) and ECD (610/20) channels, respectively. To eliminate the spillover signal of JC-1 monomers marking depolarized mitochondria into ECD channel, fluorescence compensation was performed as previously described [74].

To analyze DOXY absorption by neonatal dermal fibroblasts, NDF-3 cells were treated with different concentrations of DOXY for 72 h. DOXY fluorescence was excited by 405 nm laser and detected using the KO525 (550/40) channel [75].

### RT-qPCR

RNA isolation, reverse transcription and qPCR were performed as previously described [76]. The list of primer sequences used is provided in Supplementary Table 5. The qPCR reactions were performed in technical triplicates.

### Statistical analysis

All experiments were replicated at least three times, as detailed in the figure legends. For violin plots and bar graphs, individual data points show independent biological replicates. For all violin plots, median and quartiles are shown. Statistical analysis was performed using GraphPad Prism 8.0.2. software. Unpaired two-tailed Students’s t-test was applied when comparing 2 groups, otherwise one-way ANOVA followed by Tukey’s multiple comparison test was used. Linear correlation between 2 datasets was tested by Pearson correlation coefficient (r). p values < 0.05 were considered statistically significant; *p<0.05, **p<0.01, ***p<0.001, #p<0.0001.

## DATA AVAILABILITY

The data analyzed during this study are included in this published article and the supplemental data files. Additional supporting data are available from the corresponding author upon reasonable request.

## AUTHOR CONTRIBUTIONS

JS and KB conceived the main idea, conceptualized and drafted the manuscript; KB, MK, LYWL, PJJ, and JS designed the experiments; MDH provided neuroblastoma models and resources for validation experiments; KB and MK conducted key experiments in neuroblastoma, medulloblastoma, and schwannoma models and fibroblasts; JN, KB, and JS conducted the experiments and acquired data related to neuroblastoma tumorigenicity assays; LYWL conducted all experiments in glioblastoma and astrocytoma models; AK performed expression analysis in medulloblastoma cells; KB performed statistical analyses; KB, MK, LYWL, PJJ, MDH and JS analyzed the data; and KB, LYWL, PJJ, MDH, and JS wrote and edited the manuscript. All authors read and approved the final manuscript.

## ACKNOWLEDGEMENTS

We thank Prof. Lumír Krejčí (Department of Biology, Faculty of Medicine, Masaryk University, Brno, Czech Republic) for providing the GIMEN, LAN-1, and LAN-5 neuroblastoma cell lines, and dr. Tomáš Bárta (Department of Histology and Embryology, Faculty of Medicine, Masaryk University) for providing normal fibroblast cell lines used in this study. We also thank Johana Marešová and Jan Verner for their skillful technical assistance, and Childhood Cancer Repository (www.cccells.org) powered by Alex’s Lemonade Stand Foundation (ALSF) for tumor models.

