## Supplementary Information for "Inhibiting mitochondrial translation overcomes multidrug resistance in MYC-driven neuroblastoma via OMA1-mediated integrated stress response"

### This PDF file includes:

Legends for Supplementary Videos 1,2

Figures S1 to S11

Supplementary Tables 1 to 5

### Other Supplementary Materials for this manuscript include the following:

Supplementary Videos 1,2

Original Data with uncropped blots and replicates

**Supplementary Video 1: Mdivi-1-mediated inhibition of DRP1 induces cell death in neuroblastoma.** A time-lapse video of CHLA-15 and CHLA-20 neuroblastoma cells treated with vehicle or mdivi-1 in indicated concentrations. Phase contrast images were captured under 10× objective by the live-cell imaging system Incucyte® in 4-h intervals for 72 h.

**Supplementary Video 2: DOXY-mediated inhibition of mitochondrial translation induces cell death in neuroblastoma cells but not in nonmalignant neonatal dermal fibroblasts.** A time-lapse video of neuroblastoma cells (CHLA-15, CHLA-20) and fibroblasts (NDF-3) treated with vehicle or DOXY in indicated concentrations. Phase contrast images were captured under 10× objective by the live-cell imaging system Incucyte® in 4-h intervals for 72 h.

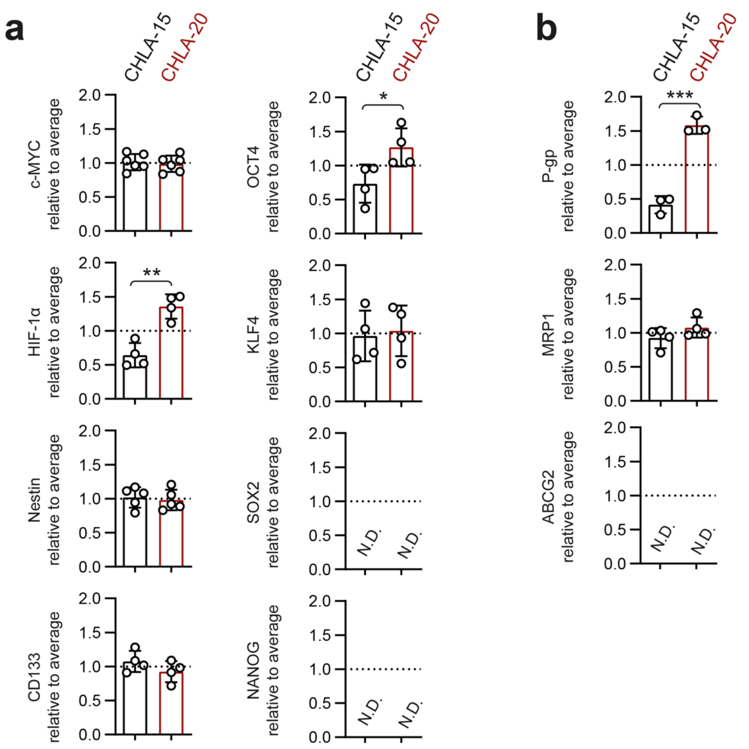


**Fig. S1**: **Analysis of cancer stemness-related markers revealed upregulation of HIF-1α and P-gp in post-therapy CHLA-20 neuroblastoma cells. a,b**, Densitometric analysis of western blotting detection of stemness transcription factors and cancer stemness-related markers (**a**) and ABC transporters (**b**) in therapy-naïve CHLA-15 and post-therapy CHLA-20 near-isogenic neuroblastoma cells. Normalized protein levels are plotted relative to the average level of both cell lines, mean ± SD; N.D. – not detected. Statistical significance was determined by unpaired two-tailed Student's t-test, **p*<0.05, **p<0.01, ***p<0.001.

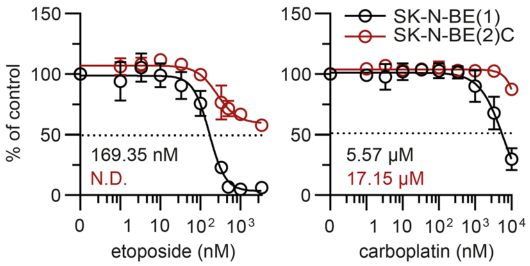


**Fig. S2: Drug sensitivity in therapy-naïve SK-N-BE(1) and post-therapy SK-N-BE(2)C near-isogenic cell lines**. CellTiter-Glo cell viability assay analysis of therapy-naïve SK-N-BE(1) and post-therapy SK-N-BE(2)C after 72-h treatment showed enhanced resistance of SK-N-BE(2)C to conventional chemotherapeutics, etoposide and carboplatin. Data are presented as mean ± SD, biological n=3, technical n=3.Calculated $IC_{50}$ are indicated. N.D.- $IC_{50}$ not determinable.

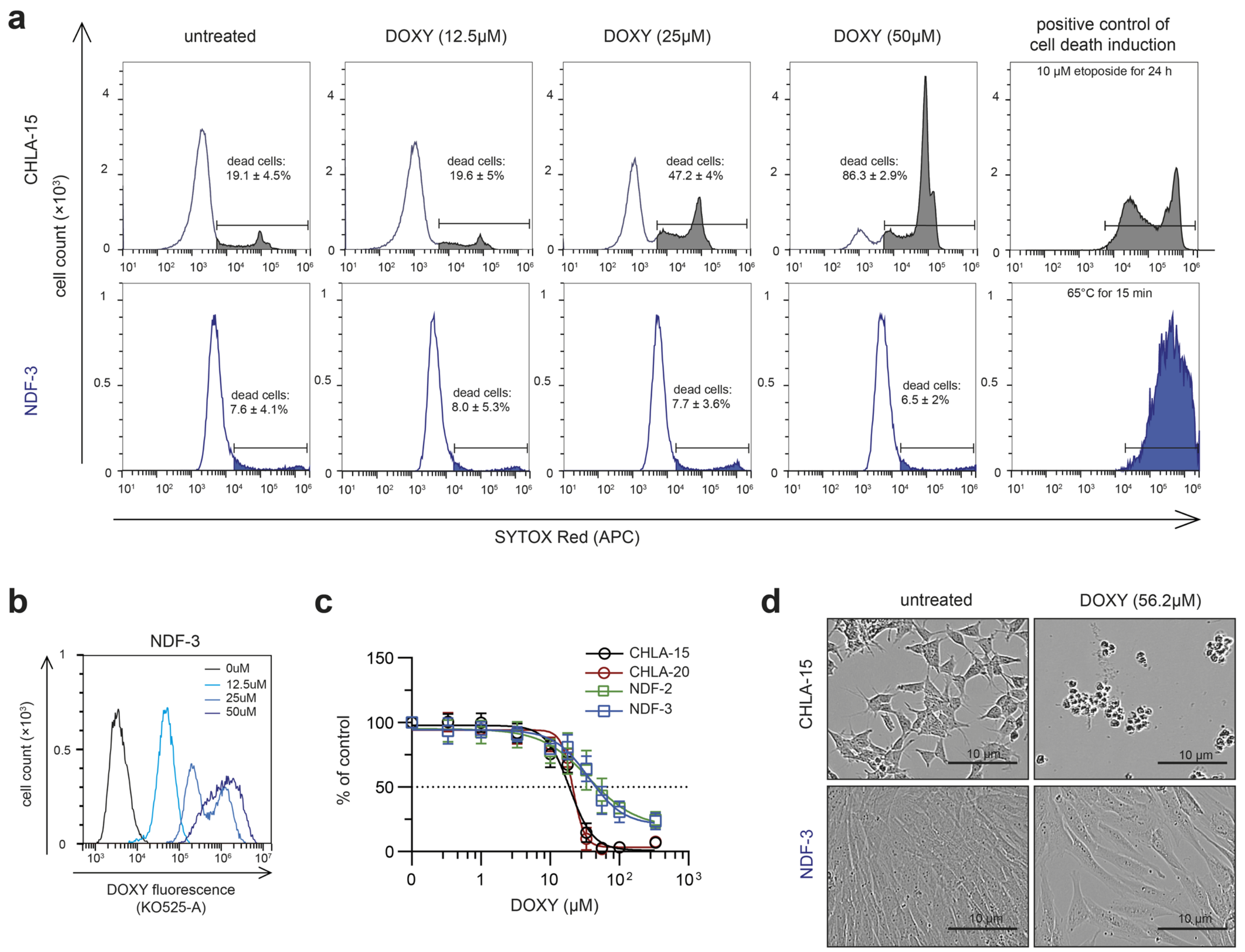


**Fig. S3**: **DOXY-mediated inhibition of mitochondrial translation does not induce cell death in neonatal fibroblasts. a**, Flow-cytometry analysis of cell viability after DOXY treatment for 72 h using SYTOX Red staining showed marked dose-dependent increase of dead cells in CHLA-15 neuroblastoma cells, whereas the viability of NDF-3 neonatal fibroblasts was not affected by the treatment. The percentages of SYTOX Red-positive dead cells are presented as mean ± SD, biological n=3**. b**, Despite having no effects on cell viability, absorption of DOXY by NDF-3 cells was confirmed by flow cytometry analysis that revealed a dose-dependent increase of DOXY fluorescence. Representative histograms are shown, biological n=2. **c,** MTT cell viability assay analysis showed that DOXY treatment limits proliferation of NDF-2 and NDF-3 neonatal dermal fibroblasts to a lesser extent compared with the CHLA-15/CHLA-20 pair. Data are presented as mean ± SD, biological n≥5, technical n≥3. The microscopic observations after 72-h treatment with indicated DOXY concentration **(d)** together with the viability analysis **(a)** indicate that DOXY-induced reduction of MTT absorbance was caused by reduction of cell growth rather than induction of cell death. See also Supplementary Video 2.

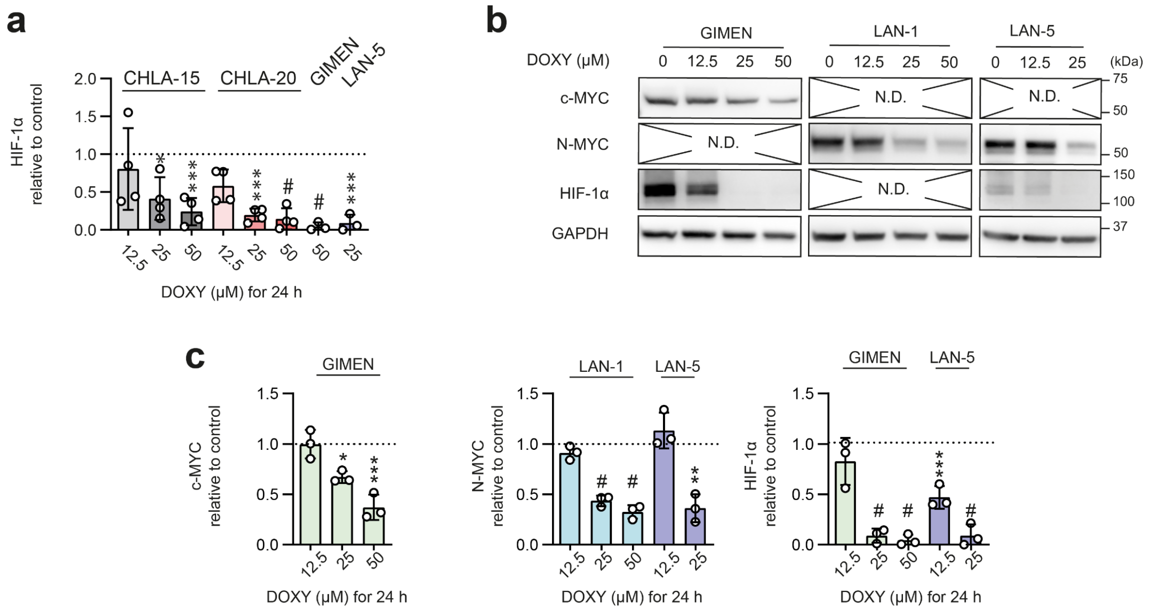


**Fig. S4: Transcription factors associated with poor neuroblastoma prognosis are downregulated upon DOXY-mediated inhibition of mitochondrial translation. a**, Densitometric analysis of western blotting detection showed significant decrease of HIF-1α across a panel of neuroblastoma cells after 24-h DOXY treatment in indicated concentrations. **b,c,** Western blotting (**b**) and densitometric analysis (**c**) showed downregulation of transcription factors c-MYC, N-MYC and HIF-1α upon 25μM and 50μM DOXY treatment for 24 h in GIMEN, LAN-1, and LAN-5 neuroblastoma cells. Normalized protein levels are plotted relative to untreated controls, mean ± SD. N.D. **-** not detectable at this cell line**.** Statistical significance was determined by one-way ANOVA followed by Tukey's multiple comparisons test (**a,c**), *p<0.05, **p<0.01, ***p<0.001, #p<0.0001.

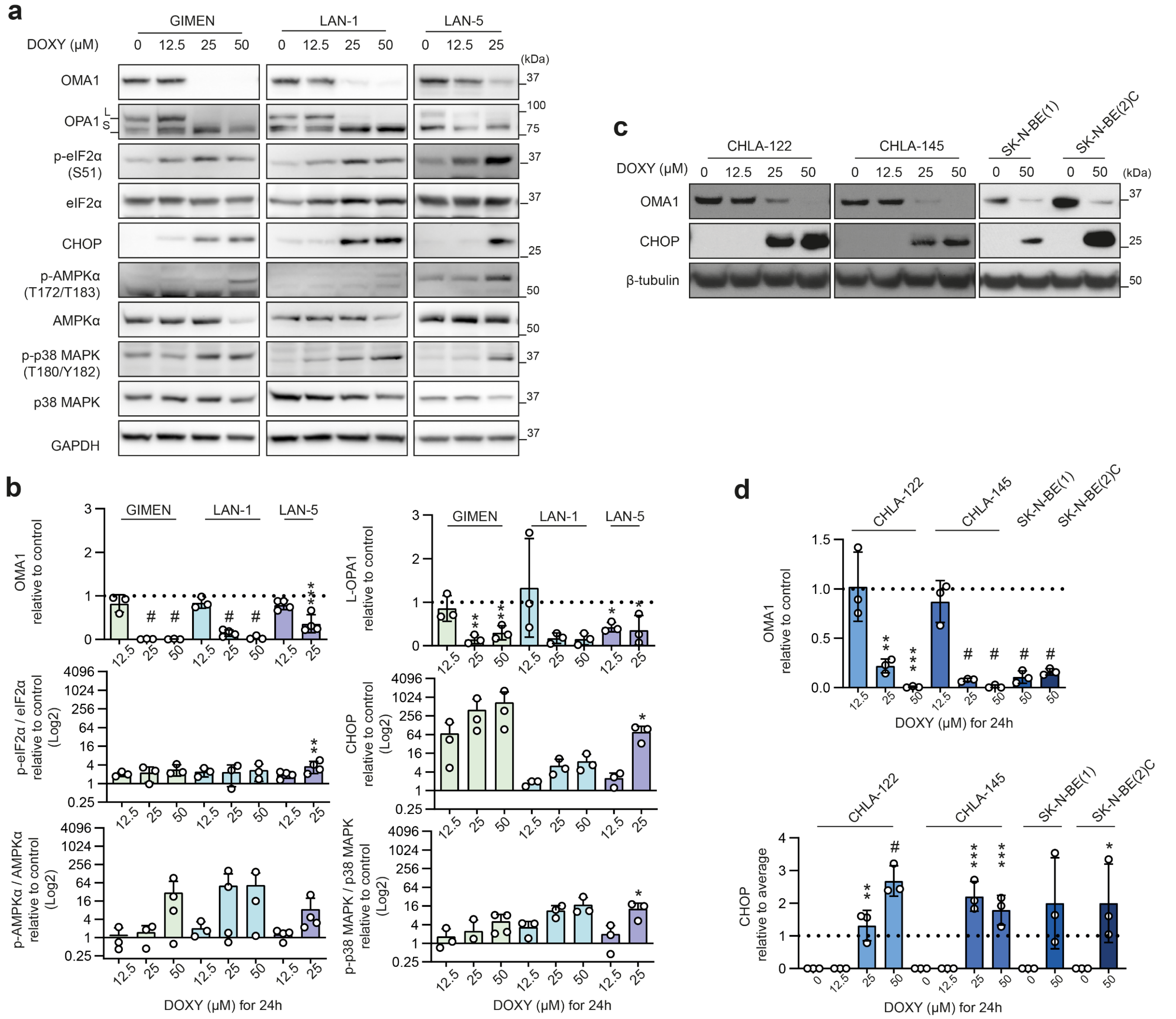


**Fig. S5: OMA1-mediated ISR is consistently activated upon DOXY-mediated inhibition of mitochondrial translation across a panel of neuroblastoma cells. a-d,** Western blotting and subsequent densitometric analysis of markers of the mitochondrial stress and ISR showed consistent effects of DOXY-mediated inhibition of mitochondrial translation across the neuroblastoma models: GIMEN, LAN-1, LAN-5 (**a,b**), CHLA-122, CHLA-145, SK-N-BE(1), SK-N-BE(2)C (**c,d**). Activating phosphorylation of AMPKα and p38 MAPK were also analyzed after 24-h treatment with indicated concentrations of DOXY in GIMEN LAN-1, LAN-5 (**a,b**). Normalized protein levels are plotted relative to untreated controls or in case of CHOP detection in CHLA-122, CHLA-145, SK-N-BE(1), SK-N-BE(2)C relative to average density of analyzed samples, mean ± SD, biological n≥3. Statistical significance was determined by one-way ANOVA followed by Tukey's multiple comparisons test (GIMEN, LAN-1, LAN-5, CHLA-122, CHLA-145) or by unpaired two-tailed Student's t-test (SK-N-BE(1), SK-N-BE(2)C), *p<0.05, **p<0.01, ***p<0.001, #p<0.0001.

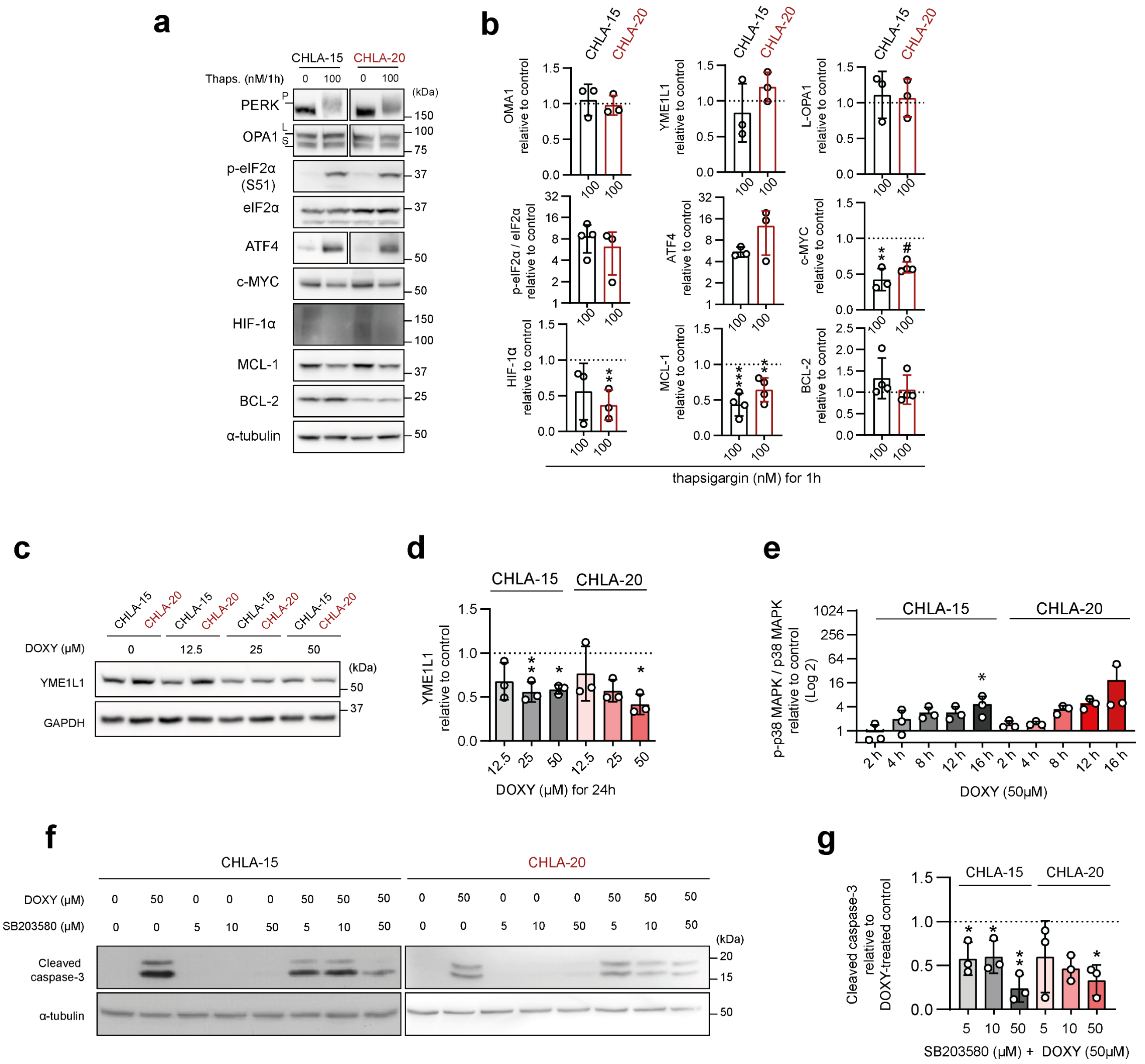


**Fig. S6: ER stress does not promote OPA1 processing and DOXY-induced cell death can be partially rescued by p38 MAPK inhibition. a,b,** Western blotting detection (**a**) and densitometric analysis (**b**) of markers of the mitochondrial stress and ISR, together with c-MYC, HIF-1α and BCL-2 anti-apoptotic proteins in neuroblastoma cells CHLA-15 and CHLA-20 after PERK-mediated activation of ISR by 100 nM thapsigargin. Note the unprocessed OPA1 and phosphorylated eIF2α in thapsigargin-treated samples. Normalized protein levels are plotted relative to untreated controls, mean ± SD, biological n=3. **c,d,** A dose-dependent downregulation of YME1L1 detected by western blotting (**c**) and densitometric analysis (**d**) in the CHLA-15/CHLA-20 pair after 24-h DOXY treatment. Normalized protein levels are plotted relative to untreated controls, mean ± SD, biological n=3**. e,** Densitometric analysis of western blotting detection revealed an increase of p-p38 MAPK (T180/Y182)/p38 MAPK in neuroblastoma cells CHLA-15 and CHLA-20 treated with 50 μM DOXY for 2–16 h. Normalized protein levels are plotted relative to untreated controls, mean ± SD. **f**,**g,** Western blotting and subsequent densitometric analysis of cleaved caspase-3 levels after 24-h 50μM DOXY treatment either w/o or with inhibitor of p38 MAPK (SB203580) showed that presence of SB203580 reduced the level of detected cleaved caspase-3 induced by DOXY comparing to DOXY alone. Normalized protein levels are plotted relative to DOXY-treated controls, mean ± SD. Statistical significance was determined by unpaired two-tailed Student's t-test (**a**) and by one-way ANOVA followed by Tukey's multiple comparisons test (**d,e,g**), *p<0.05, **p<0.01, ***p<0.001, #p<0.0001.

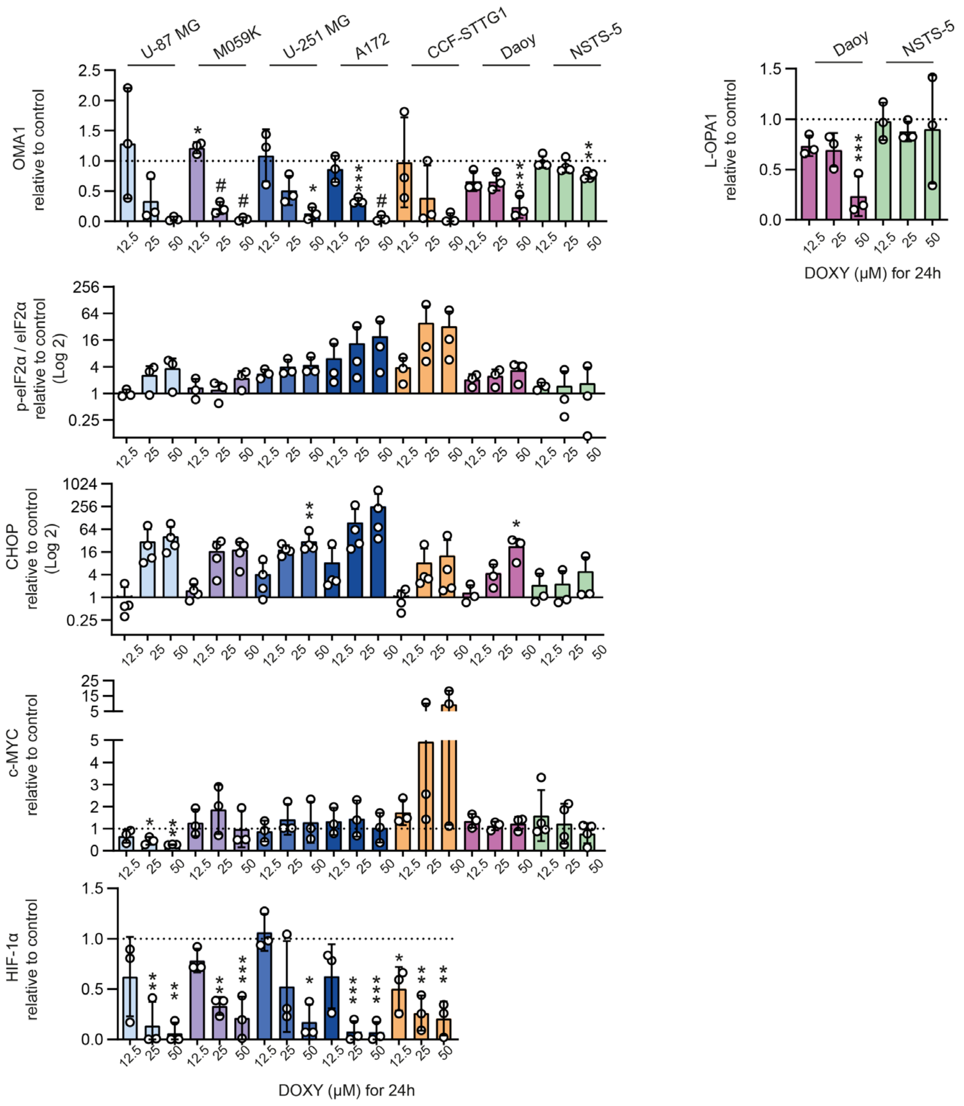


**Fig. S7: OMA1-mediated ISR is dose dependently activated by DOXY treatment in various nervous system tumors.** Densitometric analysis of western blotting detection of markers of mitochondrial stress, ISR and oncogenic transcription factors, c-MYC and HIF-1α, revealed an induction of mitochondrial stress and ISR after 24-h DOXY treatment in a panel of cell lines derived from different nervous system tumors. As in neuroblastoma, HIF-1α was significantly downregulated in nervous tissue derived tumor types, but a consistent downregulation of c-MYC was not detected. Normalized protein levels are plotted relative to untreated controls, mean ± SD. Statistical significance was determined by one-way ANOVA followed by Tukey's multiple comparisons test, *$p<0.05$, **$p<0.01$, ***$p<0.001$, #$p<0.0001$.

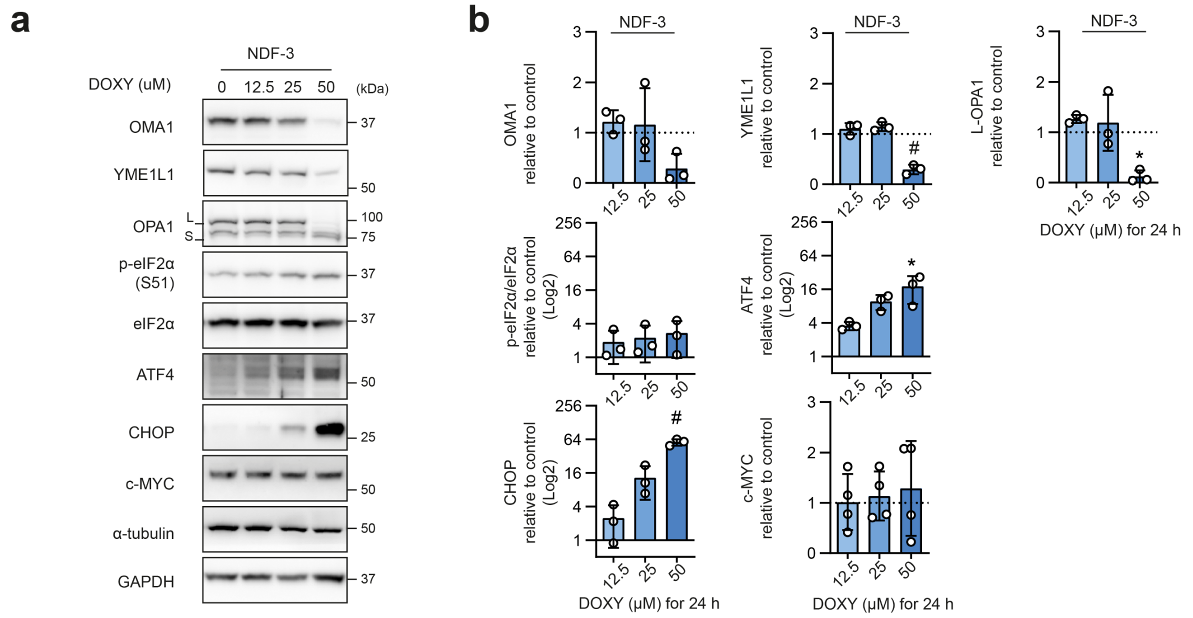


**Fig. S8: Inhibition of mitochondrial translation induces OMA1-mediated ISR in human neonatal dermal fibroblasts resistant to DOXY-induced cell death. a,b,** Western blotting detection (**a**) and densitometric analysis (**b**) showed induction of mitochondrial stress and ISR but unaffected c-MYC expression upon 24-h DOXY treatment in NDF-3 cells. Normalized protein levels are plotted relative to untreated controls, mean ± SD. Statistical significance was determined by one-way ANOVA followed by Tukey's multiple comparisons test, *$p<0.05$, #$p<0.0001$.

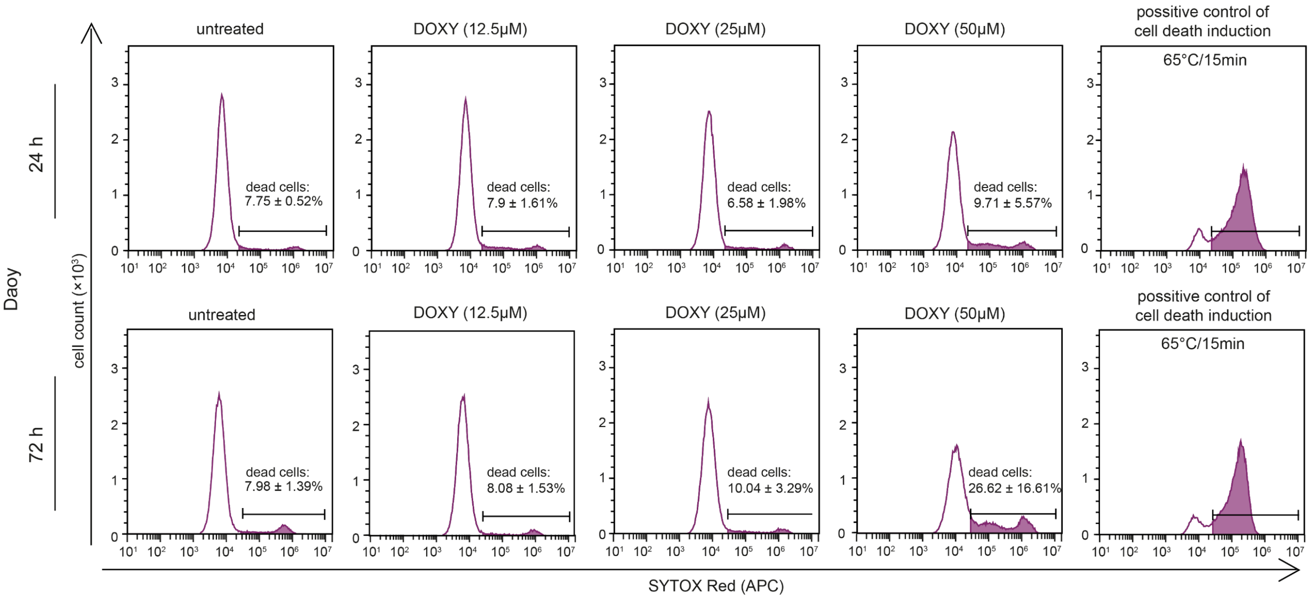


**Fig. S9: Daoy medulloblastoma cells show attenuated sensitivity to DOXY-induced cell death.** Flow cytometry analysis of cell viability using SYTOX Red staining after DOXY treatment for 24 h and 72 h. The percentages of SYTOX Red-positive dead cells are presented as mean ± SD, biological n=3 (24-h treatment) and n=8 (72-h treatment).

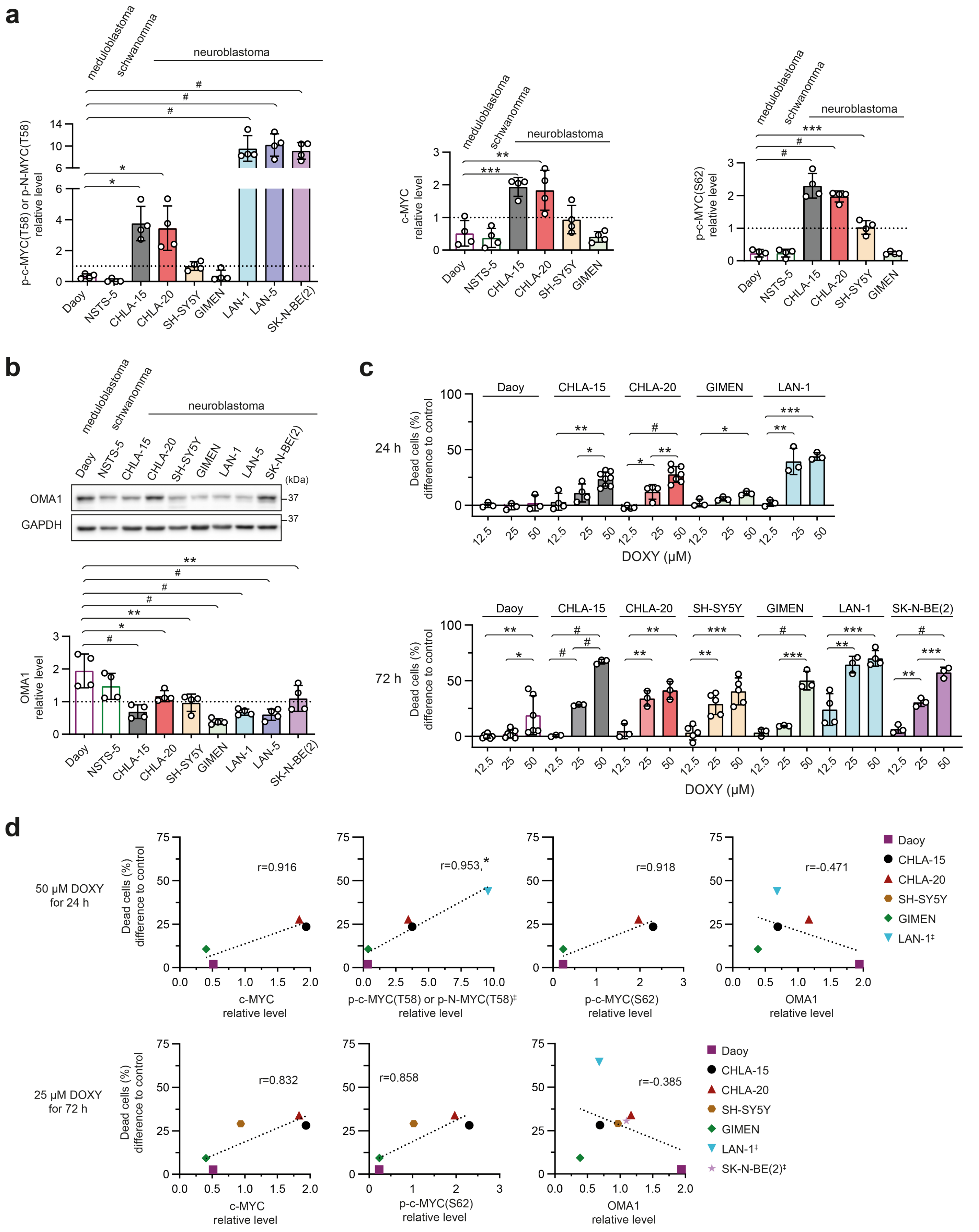

**Fig. S10: Susceptibility to DOXY-induced cell death positively correlates with overexpression and phosphorylation of MYC proteins. a-b,** Densitometric analysis of western blotting detection of p-CMYC/N-MYC(T58), c-MYC, and p-c-MYC(S62) (**a**) and OMA1 (**b**) in a panel of nervous system tumors. Normalized protein levels are plotted relative to the average level of all samples, mean ± SD. **c**, A compiled comparison of cell death rate in neuroblastoma models, CHLA-15, CHLA-20, SH-SY5Y, GIMEN, LAN-1, and SK-N-BE(2), and Daoy medulloblastoma cells after DOXY treatment for 24 h and 72 h. Viability analyzed by flow cytometry using SYTOX Red staining is presented as the difference in percentages of SYTOX Red-positive dead cells after indicated treatment vs. respective untreated controls, biological n≥3. **d**, Pearson correlation of c-MYC, p-CMYC/N-MYC (T58), p-c-MYC (S62) and OMA1 levels with difference of dead cells to control after 50 µM DOXY for 24 h and 25 µM for 72 h revealed positive correlation between high c-MYC, p-CMYC/N-MYC (T58), p-c-MYC (S62) levels and susceptibility to DOXY-induced cell death, whereas the sensitivity to DOXY was independent of OMA1 levels, r = Pearson correlation coefficient; ǂ, *MYCN*-amplified. Statistical significance was determined by one-way ANOVA followed by Tukey's multiple comparisons test (**a-c**) and by Pearson correlation (**d**), *p<0.05, **p<0.01, ***p<0.001, #p<0.0001.

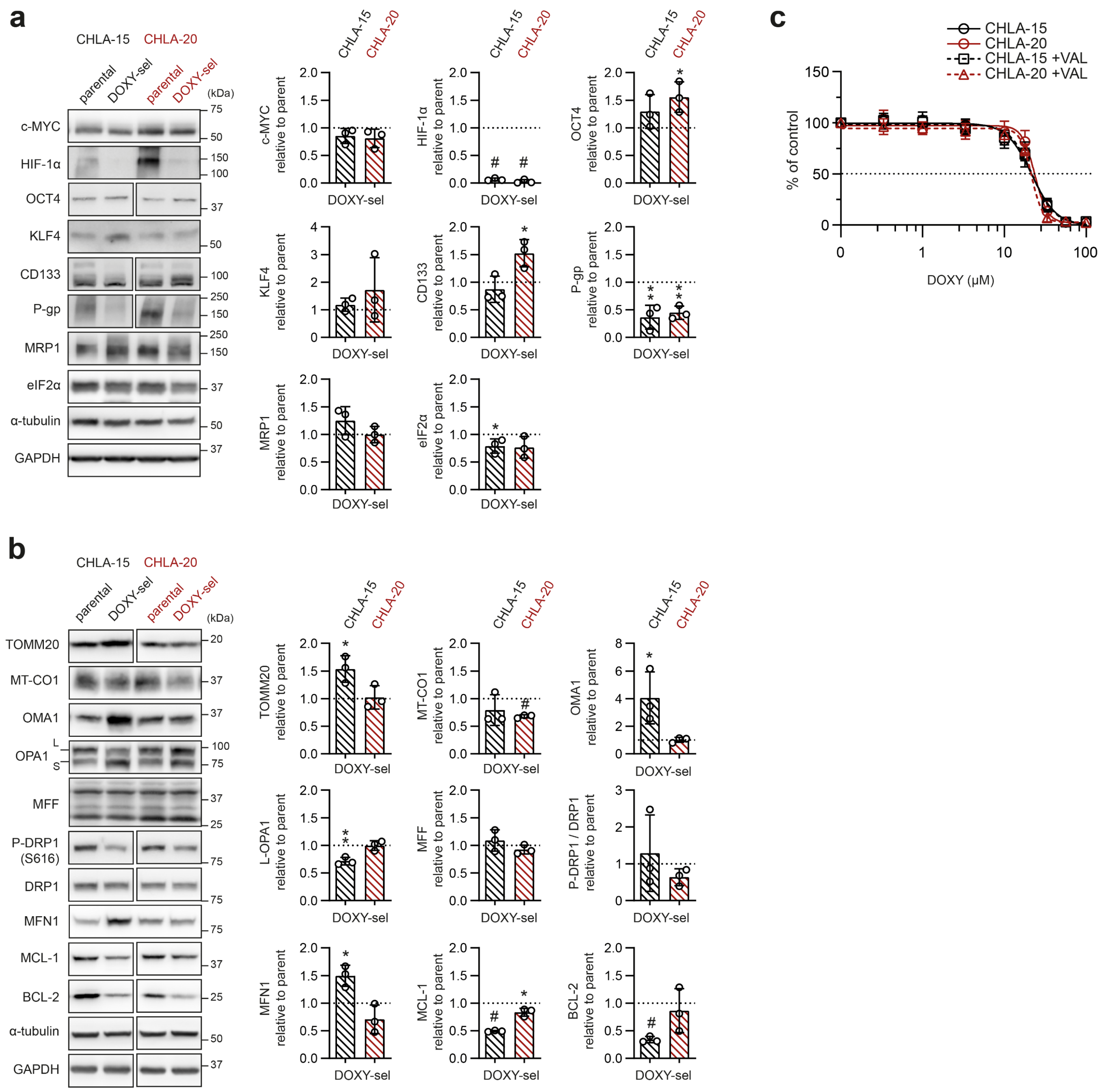


**Fig. S11: Protein expression changes in DOXY-sel cell lines. a,b,** Western blotting detection and densitometric analysis of stemness-related proteins, drug efflux pumps and eIF2α (**a**) and proteins related to mitochondrial dynamics and mitochondrial apoptotic pathway (**b**) in lysates from cells cultured in the standard medium. Normalized protein levels of DOXY-sel cells are plotted relative to normalized protein levels of parental cells, mean ± SD, biological n=3. **c,** MTT cell viability assay analysis of parent CHLA-15 and CHLA-20 cells after 72-h exposure to DOXY alone or with 0.5 μM of P-gp inhibitor valspodar (VAL). Data are presented as mean ± SD, biological n=5, technical n=3. Statistical significance was determined by unpaired two-tailed Student's t-test (**a,b**), **p*<0.05, **p<0.01, #p<0.0001.

**Supplementary Table 1. Summary of neuroblastoma cell lines and culture media composition**

| Cell Line | Cellosaurus accession # | Genetic Background | Source | Culture Medium | Supplements | Buffer System |
|---|---|---|---|---|---|---|
| CHLA-15 | CVCL_6594 | *MYC* amplified, *ALK* R1275Q | COG/ALSF Childhood Cancer Repository; www.cccells.org | DMEM/F-12 (LM-D1224, Biosera) | 10% FBS (FB-1101); 2 mM L-Glutamine (XC-T1715); 1× MEM Non-Essential Amino Acids (XC-E1154); Penicilin (100 IU/mL)/Streptomycin (100 µg/ml) (XC-A4122) all Biosera; 0.12% ITS-X (51500056, Gibco) | 15 mM HEPES & 1.2 g/l sodium bicarbonate (included in the culture medium) |
| CHLA-20 | CVCL_6602 | *MYC* amplified, *ALK* R1275Q | | | | |
| CHLA-122 | CVCL_A054 | *MYCN* amplified | | | | |
| CHLA-136 | CVCL_6590 | *MYCN* amplified | | | | |
| CHLA-145 | CVCL_AQ13 | *MYCN* amplified | | | | |
| SK-N-BE(1) | CVCL_9898 | *MYCN* amplified | | | | |
| SK-N-BE(2)C | CVCL_0529 | *MYCN* amplified | | | | |
| SH-SY5Y | CVCL_0019 | - | ECACC | | 20% FBS (FB-1101); 2 mM L-Glutamine (XC-T1715); 1× MEM Non-Essential Amino Acids (XC-E1154); Penicilin (100 IU/ml) /Streptomycin (100 µg/ml) (XC-A4122) all Biosera | |
| SK-N-BE(2) | CVCL_0528 | *MYCN* amplified | | | | |
| GIMEN | CVCL_1232 | - | Kind gift of Prof. Krejčí (Masaryk University, Brno, Czech Republic) | RPMI-1640, (PM-R1645, Biosera) | 10% FBS (FB-1101); 2 mM L-Glutamine (XC-T1715); 1× MEM Non-Essential Amino Acids (XC-E1154); Penicilin (100 IU/ml) /Streptomycin (100 µg/ml) (XC-A4122) all Biosera | 15 mM HEPES (H0887) & 1.2 g/l sodium bicarbonate (S8761), both Sigma-Aldrich |
| LAN-1 | CVCL_1827 | *MYCN* amplified | | | | |
| LAN-5 | CVCL_0389 | *MYCN* amplified | | | | |

Providers: ECACC, The European Collection of Authenticated Cell Cultures, UK Health Security Agency); Gibco (Carlsbad, CA, USA); Sigma-Aldrich (St. Louis, MO, USA). FBS, fetal bovine serum; ITS-X, Insulin-Transferrin-Selenium-Ethanolamine; HEPES, N-(2-Hydroxyethyl)piperazine-N′-(2-ethanesulfonic acid), CAS #7365-45-9.

**Supplementary Table 2. Summary of non-neuroblastoma cell lines and culture media composition**

| Cell type | Cell Line | Cellosaurus accession # | Source | Culture Medium | Supplements | Buffer System |
|---|---|---|---|---|---|---|
| medulloblastoma | Daoy | CVCL_1167 | ECACC | DMEM Low Glucose (PM-D1105, Biosera) | 10% FBS (FB-1101); 2 mM L-Glutamine (XC-T1715); 1× MEM Non-Essential Amino Acids (XC-E1154); Penicilin (100 IU/ml) /Streptomycin (100 µg/ml) (XC-A4122) all Biosera | 15 mM HEPES (H0887) & 1.2 g/l sodium bicarbonate (S8761), both Sigma-Aldrich |
| schwannoma | NSTS-5 | In-house derived (under IGA MZCR NR/9125-4 project; ethics approval no. 23/2005) | | | 20% FBS (FB-1101); 2 mM L-Glutamine (XC-T1715); 1× MEM Non-Essential Amino Acids (XC-E1154); Penicilin (100 IU/ml) /Streptomycin (100 µg/ml) (XC-A4122) all Biosera | |
| glioblastoma | U-87 MG | CVCL_0022 | ATCC | DMEM/F-12 (11320033, Thermofisher Scientific) | 10% FBS (Bovogen Biological); Penicilin (100 IU/ml)/Streptomycin (100 µg/ml) (P4333, Sigma-Aldrich) | 15 mM HEPES & 1.2 g/l sodium bicarbonate (included in the culture medium) |
| | M059K | CVCL_0401 | | | | |
| | U-251 MG | CVCL_0021 | | | | |
| | A-172 | CVCL_0131 | | | | |
| astrocytoma | CCF-STTG1 | CVCL_1118 | | | | |
| human neonatal dermal fibroblast | NDF-2 | - | Lonza Bioscience (#CC-2509), Kind gift of Dr. Bárta (Masaryk University) | DMEM High Glucose (PM-D1114, Biosera) | 10% FBS (FB-1101); 2 mM L-Glutamine (XC-T1715); 1× MEM Non-Essential Amino Acids (XC-E1154); Penicilin (100 IU/ml) /Streptomycin (100 µg/ml) (X-A4122) all Biosera | 15 mM HEPES (H0887) & 1.2 g/l sodium bicarbonate (S8761), both Sigma-Aldrich |
| | NDF-3 | | | | | |
| embryonal carcinoma | NTERA-2 clone D1 | CVCL_3407 | ECACC | | 10% FBS (FB-1101); 2 mM L-Glutamine (XC-T1715); Penicilin (100 IU/ml) /Streptomycin (100 µg/ml) (X-A4122) all Biosera | 3.7 g/l sodium bicarbonate (included in the culture medium |

Providers: ATCC, The American Type Culture Collection (Manassas, VA, USA); Biosera (Nuaille, France), Bovogen Biological (Keilor, Victoria, Australia); ECACC, The European Collection of Authenticated Cell Cultures, UK Health Security Agency); Gibco (Carlsbad, CA, USA); Lonza Bioscience (Basel, Switzerland); Sigma-Aldrich (St. Louis, MO, USA). FBS, fetal bovine serum; HEPES, N-(2-Hydroxyethyl)piperazine-N′-(2-ethanesulfonic acid), CAS #7365-45-9.

**Supplementary Table 3. Drugs used in the study**

| Drug | Manufacturer | Catalog number | Dissolvent |
|---|---|---|---|
| Ampicillin | Sigma-Aldrich | A9393 | $H_2O$ |
| ABT-737 | Abcam | ab141336 | DMSO |
| Carboplatin | EMD Millipore | 216100 | PBS |
| Chloramphenicol | Serva | 1678502 | $H_2O$ |
| Cisplatin | Sigma-Aldrich | 232120 | PBS |
| Crizotinib | Sigma-Aldrich | PZ0191 | DMSO |
| Doxorubicine | Cell Signaling Technology | 5927 | DMSO |
| Doxycycline hyclate | Sigma-Aldrich | D9891 | $H_2O$ |
| (R)-(+)-Etomoxir sodium salt | Tocris | 4539 | $H_2O$ |
| Etoposide | EMD Millipore | 341205 | DMSO |
| Linezolid | Sigma-Aldrich | PZ0014 | DMSO |
| Lorlatinib | Tocris | 5640 | DMSO |
| Mdivi 1 | Tocris | 3982 | DMSO |
| MG-132 | Tocris | 1748 | DMSO |
| Phenformin hydrochloride | Sigma-Aldrich | P7045 | $H_2O$ |
| SB203580 | Tocris | 1202 | DMSO |
| Thapsigargin | Tocris | 1138 | DMSO |
| Tigecycline hydrate | Sigma-Aldrich | PZ0021 | DMSO |
| Trametinib | APEXBIO | GEN1590424 | DMSO |
| Valspodar | Sigma-Aldrich | SML0572 | DMSO |
| Vincristine | Tocris | 1257 | DMSO |

Providers: Abcam (Cambridge, UK), APEXBIO (Houston, TX, USA), Cell Signaling Technology (Danvers, MA, USA), EMD Millipore (Billerica, MA, USA), Sigma-Aldrich (St. Louis, MO, USA), Serva (Heidelberg, Germany), Tocris (Bristol, UK).

**Supplementary Table 4. Antibodies used for immunodetection**

| Primary Antibodies (Antigen) | Manufacturer | Catalog Number | Blocking agent | Method | Dilution |
|---|---|---|---|---|---|
| ABCB1 (P-gp) | Sigma-Aldrich | P7965 | NFM | WB | 1:1 000 |
| ABCC1 (MRP1) | CST | 14685 | NFM | WB | 1:1 000 |
| ABCG2 | Abcam | ab130244 | NFM | WB | 1:1 000 |
| AMPKα | CST | 5832 | NFM | WB | 1:1 000 |
| phospho-AMPKα (Thr172/Thr183) | Abcam | ab133448 | NFM | WB | 1:1 000 |
| ATF4 | CST | 11815 | NFM | WB | 1:1 000 |
| ATP5A1 | CST | 18023 | NFM | WB | 1:2 000 |
| BCL-2 | CST | 15071 | NFM | WB | 1:2 000 |
| C-MYC | CST | 5605 | NFM | WB | 1:2 000 |
| phospho-C-MYC (Thr58) | CST | 46650 | BSA | WB | 1:1 000 |
| phospho-C-MYC (Ser62) | CST | 13748 | NFM | WB | 1:1 000 |
| CHOP | CST | 2895 | NFM | WB | 1:1 000 |
| CD133 | Abcam | ab19898 | NFM | WB | 1:1 000 |
| Cleaved Caspase 3 | CST | 9661 | BSA | IF | 1:200 |
| Cleaved Caspase 3 | CST | 9664 | NFM | WB | 1:1 000 |
| DRP1 | CST | 8570 | NFM | WB | 1:1 000 |
| phospho-DRP1 (Ser616) | CST | 4494 | NFM | WB | 1:1 000 |
| phospho-DRP1 (Ser637) | CST | 4867 | BSA | WB | 1:1 000 |
| eIF2α | CST | 5324 | NFM | WB | 1:3 000 |
| phospho-eIF2α (Ser51) | CST | 3398 | NFM | WB | 1:1 000 |
| GAPDH | CST | 2118 | NFM | WB | 1:10 000 |
| GAPDH | SCBT | sc-365062 | NFM | WB | 1:10 000 |
| HIF1α | CST | 36169 | NFM | WB | 1:1 000 |
| KLF4 | CST | 4038 | NFM | WB | 1:1 000 |
| MCL-1 | CST | 94296 | NFM | WB | 1:2 000 |
| MFF | CST | 84580 | BSA | WB | 1:2 000 |
| MFN1 | CST | 14739 | BSA | WB | 1:2 000 |
| MT-CO1 | Abcam | ab14705 | NFM | WB | 1:1 000 |
| N-MYC | CST | 84406 | NFM | WB | 1:2 000 |
| NANOG | CST | 4893 | NFM | WB | 1:1 000 |
| Nestin | Abcam | ab18102 | NFM | WB | 1:1 000 |
| OMA1 | CST | 95473 | NFM | WB | 1:1 000 |
| OPA1 | CST | 80471 | BSA | WB | 1:2 000 |
| p38αβγ MAPK | CST | 9212 | NFM | WB | 1:2 000 |
| phospho-p38 MAPK (Thr180/Tyr182) | CST | 4511 | BSA | WB | 1:1 000 |
| PERK | CST | 3192 | NFM | WB | 1:1 000 |
| PGC-1α | Abcam | ab54481 | NFM | WB | 1:1 000 |
| OCT4 | CST | 2750 | NFM | WB | 1:1 000 |
| SOX2 | CST | 3579 | NFM | WB | 1:1 000 |
| TOMM20 | SCBT | sc-17764 | BSA | IF | 1:200 |
| TOMM20 | CST | 42406 | NFM | WB | 1:3 000 |
| YME1L1 | Proteintech | 11510-1-AP | NFM | WB | 1:1 000 |
| α-tubulin | Abcam | ab7291 | NFM | WB | 1:10 000 |
| β-actin | Sigma | A1978 | NFM | WB | 1:10 000 |
| β-tubulin | CST | 86298 | NFM | WB | 1:5 000 |
| **Secondary Antibodies** | **Manufacturer** | **Catalog Number** | **Blocking agent** | **Method** | **Dilution** |
| Anti-mouse IgG, HRP-linked | CST | 7076 | NFM | WB | 1:5 000; 1:10 0000 |
| Anti-rabbit IgG, HRP-linked | CST | 7074 | NFM | WB | 1:5 000; 1:10 0000 |
| Anti-mouse IgG Alexa Fluor ® 488 | Invitrogen* | A21202 | BSA | IF | 1:200 |
| Anti-mouse IgG Alexa Fluor ® 568 | Invitrogen | A10042 | BSA | IF | 1:200 |
| **Isotype Control Antibodies** | **Manufacturer** | **Catalog Number** | **Blocking agent** | **Method** | **Dilution** |
| F (ab')2 IgG Rabbit Isotype Control | Sigma-Aldrich | ab37416-5 | BSA | IF | 1:20 000 |
| IgG2a Mouse Isotype Control | Invitrogen | MG2a00 | BSA | IF | 1:100 |

Providers: Abcam (Cambridge, UK), CST – Cell Signaling Technology, Inc.,(Danvers, MA, USA), Invitrogen, (Carlsbad, CA, USA), Proteintech Group (Rosemont, IL, USA), SCBT – Santa Cruz Biotechnology, Inc., (Dallas, TX, USA), Sigma-Aldrich (St. Louis, MO, USA). BSA, bovine serum albumin; IF, immunofluorescence staining; NFM, dry non-fat milk; WB, western blotting.

**Supplementary Table 5. The sequences of primers used for RT-qPCR**

| List of primers | | |
|---|---|---|
| **Gene** | **Forward (5'→3')** | **Reverse (3'→5')** |
| *HIF1A* | CAAGAACCTACTGCTAATGC | TTATGTATGTGGGTAGGAGATG |
| *HSP90AB1* | CGCATGAAGGAGACACAGAA | TCCCATCAAATTCCTTGAGC |
| *MCL1* | GGACAAAACGGGACTGGCT | ATGCCACCTTCTAGGTCCTCT |
| *MYC* | TCTCTCCGTCCTCGGATTCT | GCCTCTTTTCCACAGAAACAACA |
